# Sparse input representations explain odor discrimination in complex, concentration-varying mixtures

**DOI:** 10.64898/2026.01.27.702074

**Authors:** Hannah McCalmon, George Cai, Constantine Tsibouris, Noé Hamou, Jo Hoskins, Farhad Pashakhanloo, SueYeon Chung, Vikrant Kapoor, Venkatesh N. Murthy

## Abstract

In natural environments, animals must detect behaviorally relevant odors despite variability in both odor mixture composition and stimulus intensity. Although mice can identify salient odors embedded in complex mixtures, how target concentration and background complexity jointly constrain discrimination remains unclear. We trained mice in a two-alternative forced choice task to identify target odors embedded in mixtures containing up to 16 background components. After performance stabilized, we systematically varied target odor concentration. Discrimination accuracy declined with decreasing target concentration but showed little additional dependence on background complexity. Using a biophysically grounded model of olfactory bulb glomerular responses, we show that linear decoding reproduces behavioral performance when intrinsic neural noise dominates over background-driven variability. Manifold capacity analysis revealed that neural representations remain efficiently structured for odor discrimination despite variation in background complexity. These results define a noise-limited regime of olfactory discrimination in which target detectability is primarily constrained by neural sensitivity rather than background interference.

## INTRODUCTION

Animals rely on sensory inputs to identify food^1,2^, locate mates^3^, avoid predators^4,5^, and detect environmental hazards^6,7^. In natural environments, these signals are embedded within complex and dynamic sensory scenes, where relevant cues must be extracted from competing background inputs. The process of scene analysis enables the brain to filter and organize sensory information, enhancing salient signals while suppressing irrelevant or interfering background activity^8,9^. This challenge is conserved across modalities. In audition, it is exemplified by the “cocktail party effect,” where a single voice can be isolated amid background noise^10,11^, and in vision, by figure-ground segmentation in cluttered environments^12,13^. Across these systems, perceptual limits often reflect a balance between the strength of relevant signals and interference from background inputs.

Whether the olfactory system supports analogous forms of scene analysis, and what factors limit this ability, remains unclear^14–18^. In recent work, we demonstrated that mice can detect behaviorally relevant target odors embedded in complex, variable backgrounds^19^. However, performance deteriorates when background odors activate overlapping glomeruli in the olfactory bulb, suggesting that interference at the level of population representations can obscure target signals. The degree of interference can also depend on the relationship between target and background odors. In both humans and rodents, backgrounds that are perceptually or neurally similar to the target can impair detection more strongly than dissimilar backgrounds, suggesting that target-background congruence is an important determinant of figure-ground segregation^19–21^. Computational analyses have shown that odor representations can remain linearly separable at the population level under certain conditions^22^, and humans are similarly capable of identifying individual mixture components^23^. These findings raise the question of what ultimately limits mixture discrimination: interference from overlapping background activity, insufficient signal strength, or an interaction between the two.

A key variable that may govern this balance is odor concentration, as it relates to the sensitivity of the receptor repertoire. In natural environments, odor signals exhibit large fluctuations in concentration due to turbulent transport in airborne plumes^24–26^, as well as factors such as volatility and substrate disruption in surface-deposited cues. In addition, respiratory dynamics, including sniff frequency and inhalation volume, modulate the number of odorant molecules reaching the epithelium^27–29^. Together, these factors can strongly influence the strength and reliability of sensory input, effectively placing the system in different noise regimes. Despite this, how variations in signal strength interact with background interference to shape perception remains poorly understood.

Here, we ask how target odor concentration and background complexity jointly determine performance in an odor-guided discrimination task. Specifically, we test whether limits on mixture discrimination arise primarily from weak sensory signals, from interference due to overlapping background activity, or from their interaction. We trained mice to identify target odors embedded in mixtures containing variable numbers of background components while systematically varying target concentration over several orders of magnitude. To interpret these behavioral results, we analyzed a biophysically grounded model of olfactory bulb glomerular activity, allowing us to examine how changes in signal strength and background structure shape the geometry and linear separability of population-level odor representations. Together, this approach provides a unified framework for understanding how neural noise and background interference constrain olfactory scene analysis.

## RESULTS

### Learning stabilizes performance across odor background complexity

We trained mice to report the identity of one of two target odors by licking the corresponding water spout (Figure 1A,S1). Trials were self-initiated upon entry into the odor port, triggering a 1-s odor presentation. Correct responses were rewarded with water, while incorrect choices led to reward omission. Trials ended at the first lick or after 20 s.

**Figure 1.**
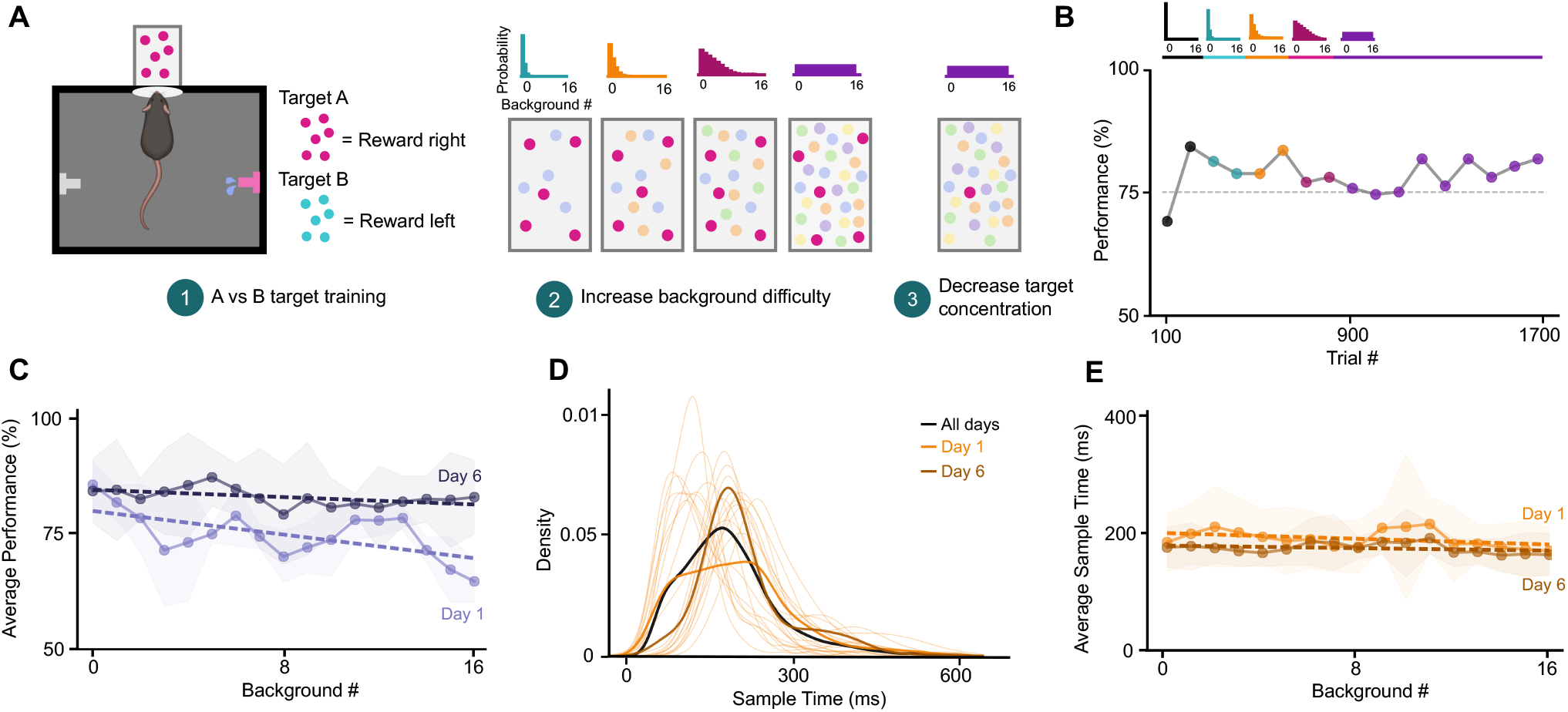
Performance improves with training across increasingly complex background conditions. (A) Schematic of training structure in a freely moving 2AFC odor discrimination task. Dots represent individual odorant molecules with different colored dots representing different odors. (B) Average performance across increasing background complexities. Trials are binned by 100. All background complexities besides the most complex, uniform probability distribution, represent the first and last 100 trials for the given condition. Gray dashed line indicates criterion performance (75%). (C) Performance as a function of background complexity for Day 1 and Day 6 (two-way ANOVA, *p* = 5.8 *×*10^*−*5^). Dashed lines indicate linear fit, showing reduced impact of background complexity over learning. Shaded region represents ± SEM across animals. (D) Trial-wise distributions of sample times at the highest background complexity. Thin lines show individual-day distributions, the bold black curve shows the overall average, and light versus dark curves indicate Day 1 and Day 6 of training. (E) Sample time showed did not change across background complexity from Day 1 to Day 6 (two-way ANOVA, *p* = 0.081). Shaded region represents ± SEM across animals. Data are from *n* = 4 animals.

After initial training on the two targets, mice were presented with the same targets embedded in mixtures of randomly selected background odors. During all trials, the concentration of the individual background odors remained constant (5 % (v/v)). The number of background components per trial was gradually increased until it approximated a uniform distribution across 0–16 components once animals reached criterion performance (75%).

Under this uniform mixture distribution, mice reached criterion performance rapidly and continued to improve over approximately 1,000 trials (Figure 1B). Early in training, performance declined with increasing background complexity. This dependence diminished with continued training, and by Day 6, performance became largely independent of background complexity (Figure 1C). A two-way ANOVA confirmed significant effects of background complexity and training day, as well as an interaction between them, indicating that the impact of background complexity decreased with learning.

To determine whether these improvements reflected changes in behavioral strategy, we examined sampling and action times. Neither sample time nor action time varied systematically with background complexity or across training (all *p* > 0.05; Figures 1D,E,S2A-C). Thus, improvements in performance cannot be explained by changes in sampling duration or motor execution.

Together, these results show that with experience, mice reach a stable behavioral regime in which performance is largely independent of background complexity and not accounted for by changes in sampling or action timing. With a consistent behavioral baseline, we could next ask how signal strength and background interference limit mixture discrimination.

### Performance degrades systematically with decreasing target odor concentration

Once mice achieved criterion performance under the uniform background distribution, we systematically reduced the target odor concentration. Mice were initially trained at either 5 % or 2.5 % (v/v), and concentrations were serially diluted (5.0, 2.5, 0.5, 0.25, 0.05, 0.005 %). Animals spent on average 4.19 ± 0.57 days (3949 ± 372 trials) per condition. As target odor concentration decreased, performance declined systematically, following a sigmoidal relationship (Figure 2A). A logistic function provided a good descriptive fit to the aggregated performance data (*R*^2^ = 0.58).

**Figure 2.**
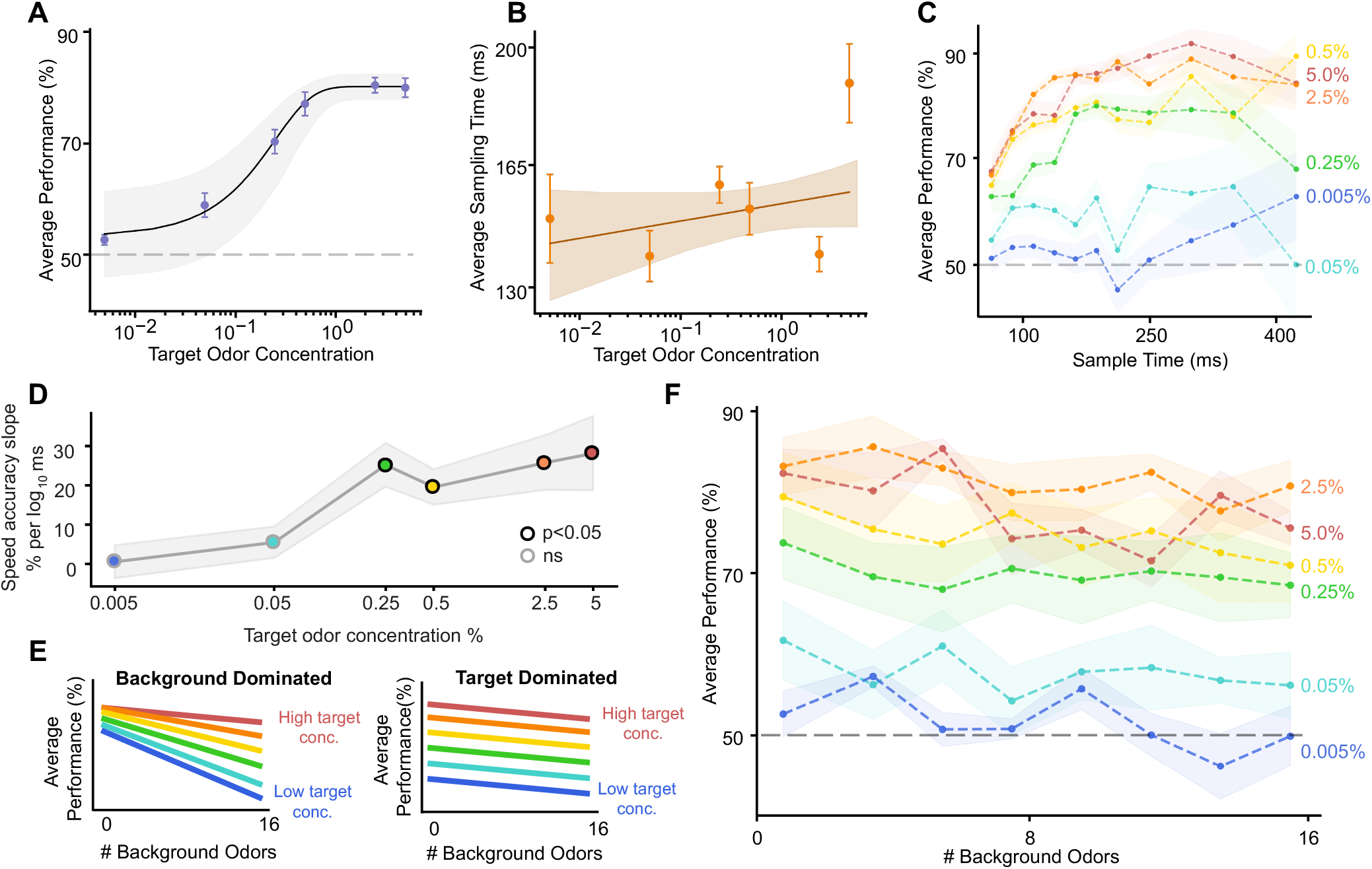
Performance depends on target odor concentration, not background complexity. (A) Average performance increased with target odor concentration, following a sigmoidal trend. Points show mean success rate across mice and training days at each concentration; error bars indicate ±SEM. A logistic function provided a good descriptive fit to the aggregated data (solid curve; *R*^2^ = 0.58), with shaded area indicating the 95% confidence interval. The dashed line denotes chance performance (50%). Performance increased significantly with concentration (trial-level mixed-effects logistic regression with animal as a random effect, *p <* 0.001). (B) Sample time increased modestly with concentration (log-linear regression, *p <* 0.05). Points show mean sample time across all mice and training days. Error bars show ±SEM across mice and days at each concentration; orange shading indicates the 95% confidence interval of the loglinear fit. (C) Across concentrations, longer sampling improved performance primarily at higher concentrations (two-way ANOVA, interaction *p <* 0.05). (D) For each concentration, a binomial logistic regression was fitted to the relationship between the performance and the odor port sampling time (log_10_), using binned trial counts as the response variable. The speed-accuracy slope (y-axis) depicts the estimated change in performance (% correct) per log_10_ unit increase in sample time, calculated via parametric bootstrap. Points show the mean slope; shaded region indicates ±SE from bootstrap. Black circles indicate concentrations at which the slope was significantly greater than zero (*p <* 0.05); gray circles indicate non-significant slopes. (E) Conceptual illustration of two models for how background complexity could influence performance. In a background-dominated model, performance decreases with increasing background odors; in a target-dominated model, performance scales with target concentration regardless of background complexity. (F) Performance across background complexities at six target odor concentrations spanning three orders of magnitude. Dashed lines show group means and shaded areas denote ±SEM across animals. Performance increased significantly with concentration (one-way ANOVA, *p <* 0.001) but showed no systematic effect of background complexity. Data are from *n* = 6 animals.

To assess the effect of concentration while accounting for repeated measurements within animals, we fit a trial-level mixed-effects logistic regression with animal as a random effect. Increasing concentration significantly increased performance 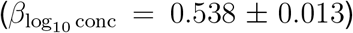, corresponding to a 1.71-fold increase in the odds of a correct response for each 10-fold increase in concentration (95% CI: 1.67–1.76; *p* < 10^−300^; *n* = 6 animals, 22,173 trials).

Further analysis confirmed that the variation of performance across target odor concentrations was consistent across animals and training stage (Figure S3A–D). When low-concentration targets were presented unpredictably as interleaved “probe” trials rather than through continual training, performance declined more sharply with lower target concentration, indicating reduced sensitivity without repeated experience at those weak concentrations (Figure S3C). Together,these analyses reveal a robust, experience-dependent effect of target odor strength on discrimination accuracy.

To determine whether these performance changes reflected altered decision strategy or differences in sensory evidence, we examined trial timing. Sampling times decreased modestly with concentration (log-linear regression, *p* = 0.018; Figure 2B), indicating that animals made faster, less accurate choices when sensory evidence was weak, consistent with previous observations in olfactory decision-making^30^. Action times showed a similar, non-significant trend (log-linear regression, *p* = 0.086; Figure S2C).

Critically, the benefit of extended sampling depended on odor strength. At high concentrations, performance increased with longer sampling durations (Figure 2C), consistent with a classical speed-accuracy trade-off^31,32^, whereas this relationship was absent at low concentrations. To further characterize how the benefit of extended sampling depended on odor strength, we estimated the speed-accuracy slope (the rate at which performance improved with sampling duration) separately at each target concentration (Figure 2D). At high concentrations (0.25%–5%), the slope was significantly positive (binomial generalized linear model, GLM, *p* < 10^−8^ at all four concentrations), indicating that longer sampling improved discrimination. In contrast, the slope was near zero and non-significant at the two lowest concentrations (0.05%: slope = 5.4%, *p*= 0.21; 0.005%: slope = 0.5%, *p* = 0.91), indicating that additional sampling provided no benefit when the target signal was weakest. The transition between these two regimes occurred between 0.05% and 0.25%, closely matching the inflection point of the psychometric curve (Figure 2A). These results indicate that the speed-accuracy tradeoff is concentration-dependent: at high concentrations performance is limited by evidence accumulation and can be improved by longer sampling, whereas at low concentrations performance is constrained by signal strength and additional sampling is uninformative.

We next asked whether the decline in performance at low concentrations reflected increased interference from background odors or a reduction in target signal strength. In a background-dominated regime, weak targets would be increasingly masked by distracting odors, leading to a strong dependence of performance on background complexity. In contrast, in a target-dominated regime, performance would decline uniformly across background conditions as target concentration decreased (Figure 2E). The data support a target-dominated regime: performance declined systematically with decreasing target odor concentration and showed only a weak additional dependence on background complexity (Figure 2F). A one-way ANOVA across concentrations revealed a highly significant main effect of concentration (*F*_5, obs_ = 12.56, *p* = 1.3 × 10^−6^) but no additional effect of background complexity. Tukey HSD post-hoc comparisons showed that discrimination accuracy at the weakest target odor concentration (0.005%) was significantly lower than at higher concentrations (0.25–5.0;% *p* < 0.01), while differences among higher concentrations were not significant. All behavioral effects were replicated using a distinct target-odor pair, demonstrating that these phenomena generalize across odor identities (Figure S4).

Together, these results show that target odor concentration is the primary factor limiting performance, with background complexity playing a comparatively minor role. As target signals weaken, discrimination accuracy declines in a manner largely independent of background interference, consistent with a noise-limited regime. Behavioral adjustments in sampling track stimulus strength but do not account for the observed performance deficits, indicating that limits on mixture discrimination arise from the strength of the sensory signal itself.

### Modeling Mouse Performance with Linear Decoders

We next asked whether empirically observed early olfactory neural representations are sufficient to account for the behavioral pattern observed in our task via linear decoding of glomerular responses^22^. We use this model to provide a rationalization for how the glomerular representation might support or limit odor discrimination performance, though the modeled neural representation was static. Behaviorally, performance declined at low target odor concentrations not because mice selectively failed in complex mixtures, but because performance decreased similarly across background conditions (Figure 2F). This suggested a target-dominated regime in which performance was primarily limited by the detectability of weak target signals rather than by interference from background odors.

To test how such a regime could arise, we trained linear decoders on a biophysically motivated model of olfactory bulb glomerular responses^33^ (Figure 3A, model described in section Methods: Biophysical Model). This framework allowed us to independently manipulate neural noise, receptor interactions, and the concentration and complexity of background mixtures to determine which factors best accounted for the behavioral results.

**Figure 3.**
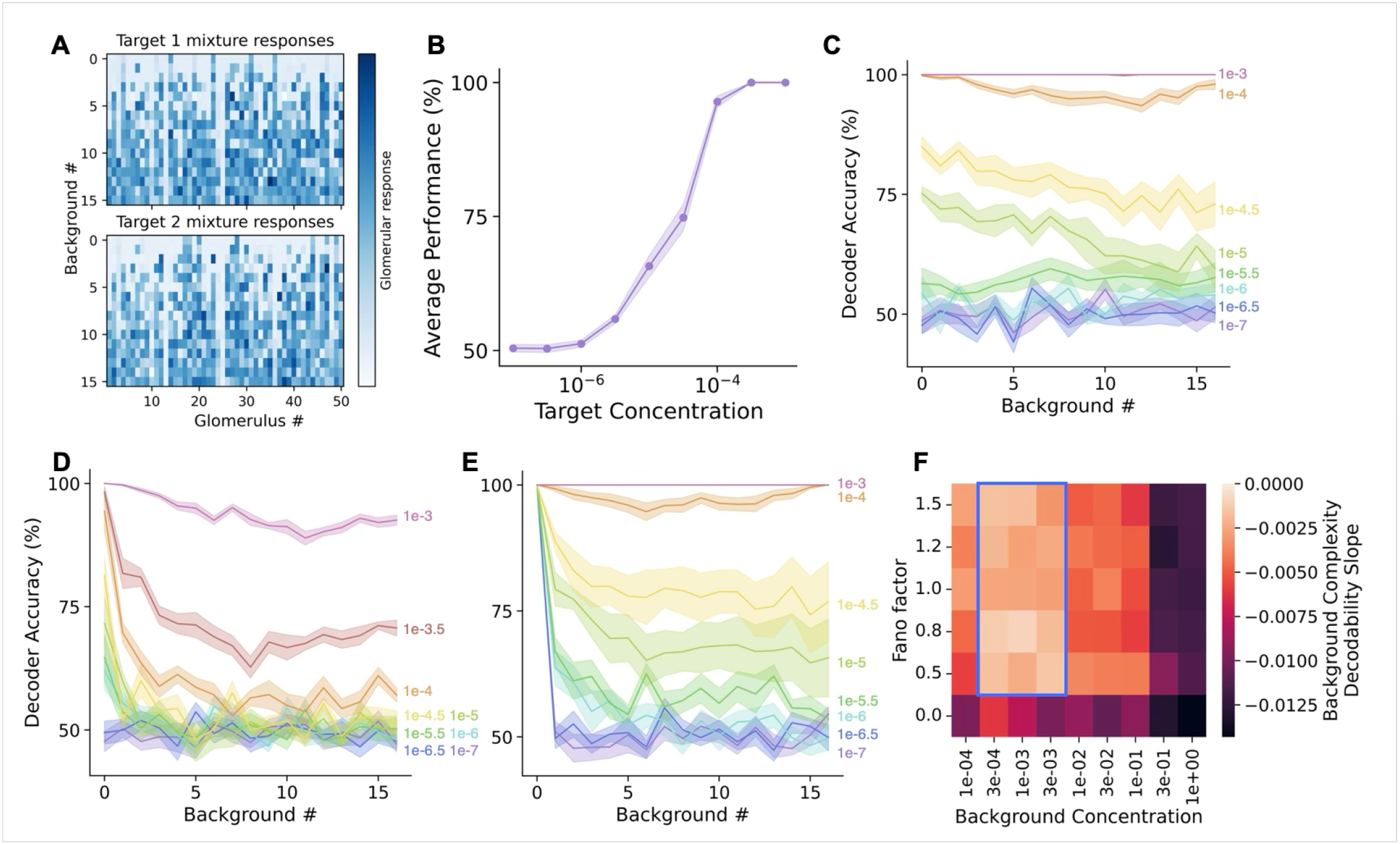
Behavioral performance may be explained by relative strengths of neural and background noise. (A) Example glomerular model responses to mixtures with different numbers of background odors. The first 16 rows are mixtures containing target 1, and the last 16 rows contain target 2. The target and background concentration used was 10^*−*2^ and 10^*−*2.5^ respectively, which we designate as “weak.” (B) Decoder performance averaged across background mixtures as a function of target concentration. The target concentration during training was 10^*−*2^ and background concentration was 10^*−*2.5^. (C) Decoding accuracy curves with Poisson-like neural noise and weak training target and background odor concentration (10^*−*2^ and 10^*−*2.5^ respectively). Solid line indicates mean decoding accuracy; shaded region indicates SEM across 10 replicate datasets. (D) Decoding accuracy curves with Poisson-like neural noise. The target concentration during training was 10^*−*2^ while background odors were at a “strong” concentration 10^*−*1^. (E) Decoding accuracy curves with neural noise set to 0 using the same concentrations as in (C). (F) Average slope of background odors against decoder accuracy as a function of background concentration and neural noise strength as quantified by the Fano Factor. Train target concentration was weak (10^*−*2^). Note that the steepest possible slope of *−*0.03125 corresponds to perfect performance with 0 background odors and chance level performance with 16 background odors.

We first examined whether decoder performance was more sensitive to target concentration or to background mixture complexity. Decoder performance declined systematically as target concentration decreased (Figure 3B), recapitulating the concentration dependence observed behaviorally (Figure 2A). Under weak background conditions, increasing mixture complexity had little effect on decoding performance (Figure 3C), consistent with the target-dominated regime observed in behavior. By contrast, when background concentrations were high, increasing mixture complexity substantially impaired decoding as target concentration decreased (Figure 3D), demonstrating that a background-limited regime can emerge under sufficiently strong mixture concentrations.

We next asked what factors limited performance in the weak background regime that best matched behavior. In the absence of Poisson-like neural noise, decoder performance was largely preserved except under the conditions with few (0-3) background odors (Figure 3E). Consistent with this interpretation, the weak-background regime quantitatively matched behavioral performance substantially better than the strong-background regime (RMSE = 0.118 vs. 0.245; MAE = 0.099 vs. 0.223 paired Wilcoxon signed-rank test on absolute errors, *p* = 0.042) and parameter sweeps across background concentration and neural noise similarly identified weak-background conditions with Poisson-like neural variability as the regime most consistent with behavior (Figure 3F). Together, these results indicate that behavioral performance declines because neural variability masks weak target signals before background mixtures become strongly interfering, consistent with a noise-limited rather than background-limited regime.

Having identified neural variability as the dominant limitation under our task conditions, we next asked whether this interpretation depended strongly on assumptions about receptor-level mixture interactions. Odor receptor responses to complex mixtures are not linearly additive, and odors show antagonistic interactions, effectively normalizing the receptor population response and decreasing interference from background odors^33,34^. Antagonistic interactions were present in the model by default, but removing receptor antagonism had little effect on the weak-background regime that best matched behavior, suggesting that the core result does not depend sensitively on the precise form of receptor interactions (Figure S5A).

Although the sensory regime was best explained by neural variability, another factor that could influence behavioral performance is how the decision boundary is updated as target concentrations decline. In our initial modeling framework, decoders were trained only at high target concentrations, implicitly assuming that the same readout could be applied unchanged across dilutions. However, mice continued learning throughout the target dilution sessions, potentially adapting the decision boundary over time. When we trained and tested decoders at matched target concentrations, performance improved substantially relative to the fixed-decoder case, but no longer matched the behavioral performance curves observed in mice (Figure S5C). Training decoders across the full range of target concentrations produced similar results (Figure S5D), indicating that learning to separate a pair of targets across a wide range of concentrations simultaneously is possible. Together, these results suggest that mice partially update the decision boundary as target concentrations decline, consistent with the improved performance observed during continual target dilution training relative to randomly interleaved probe trials (Figure S3C).

Across these analyses, weak target signals consistently became limiting before background mixtures became strongly interfering, despite variations in receptor interactions and decoder training conditions. We therefore next asked what features of the sensory representation could support such robustness to background odors. Decoder performance depended strongly on the structure of the readout: sparse readouts (L1 regularization; Figure S5A), which rely on a relatively small subset of glomerular inputs, better reproduced the behavioral regime than dense readouts (L2 regularization; Figure S5B), which pool information across many glomeruli. Although this difference was initially unexpected, sparse readouts are biologically plausible given that downstream piriform cortical neurons do not form all-to-all connections with olfactory bulb inputs^35–37^.

This sparse-coding interpretation also provides a parsimonious explanation for why background mixtures produced relatively little interference even as target concentrations declined. Sparse glomerular representations reduce cross-talk between target and background odors, limiting mixture interference even in complex odor environments. Consistent with this interpretation, the sparsity distribution produced by our weak-background regime (10^−2.5^) fell between two published datasets measuring glomerular responses to monomolecular odors at relatively high and minimal concentrations, respectively^34,38^ (Figure S6A).

To assess whether target-background similarity could account for the concentration-dependent performance decline, we quantified glomerular masking similar to Rokni et al. (2014)^19^ using published imaging data (Zak et al. 2020)^34^ and our biophysical model (Figure S6B-D). Masking increased with background complexity and depended on target identity, with structurally similar backgrounds contributing more to interference. However, masking changed only weakly across target concentrations despite behavioral performance declining steeply over the same range. This dissociation indicates that target-background masking alone cannot explain the concentration-dependent performance changes and is consistent with our conclusion that performance in this task is primarily constrained by neural noise rather than background interference.

Taken together, behavior, modeling, and physiology converge on the conclusion that neural variability, rather than background mixture complexity, is the principal limitation under our task conditions. Our results further suggest that sparse sensory representations help preserve robustness to background odors even as target signals weaken. The model predicts that only substantially (> 1 order of magnitude) stronger backgrounds or much more complex mixtures would shift performance into a truly background-limited regime (Figure 3E), a hypothesis that can be tested experimentally.

### Analyzing the Geometry of Glomerular Neural Manifolds

The previous analyses established that neural variability and sparse coding strongly affect linear decodability, but they do not address how these factors alter the structure of the underlying glomerular representations. In particular, it remained unclear whether weak target signals primarily reduce separability by increasing the magnitude of variability within each target representation, by increasing the number of dimensions along which the representation varies, or both. To address this, we used the neural manifold capacity framework^39–41^, which provides a geometric description of how easily neural representations can be separated.

Conceptually, the point cloud of population responses to all odor mixtures containing a given target odor comprises a neural manifold. The framework quantifies both how much variability exists within a manifold and how easily different manifolds can be separated. Variability within a manifold is characterized by its effective dimensionality and effective radius, which respectively measure the number of directions along which the representation varies and the magnitude of that variability. Together, these geometric statistics determine the manifold capacity, a measure of how easily two or more neural manifolds can be linearly separated (Figure S8).

Using this framework, we examined how neural variability and background complexity jointly shaped glomerular representation geometry. The primary measure in manifold analysis is how easily the two cue representations can be linearly separated, quantified by the manifold capacity. Increasing neural noise (black dashed line) consistently reduced manifold capacity, indicating that the cue manifolds became progressively less separable in neural activity space (Figure 4). The framework further measures how the linear separability arises due to variability within each cue manifold through the effective dimension and radius. Increasing neural noise similarly increased effective dimension and radius, indicating that noise in the glomerular response globally broadened and destabilized the cue representations.

**Figure 4.**
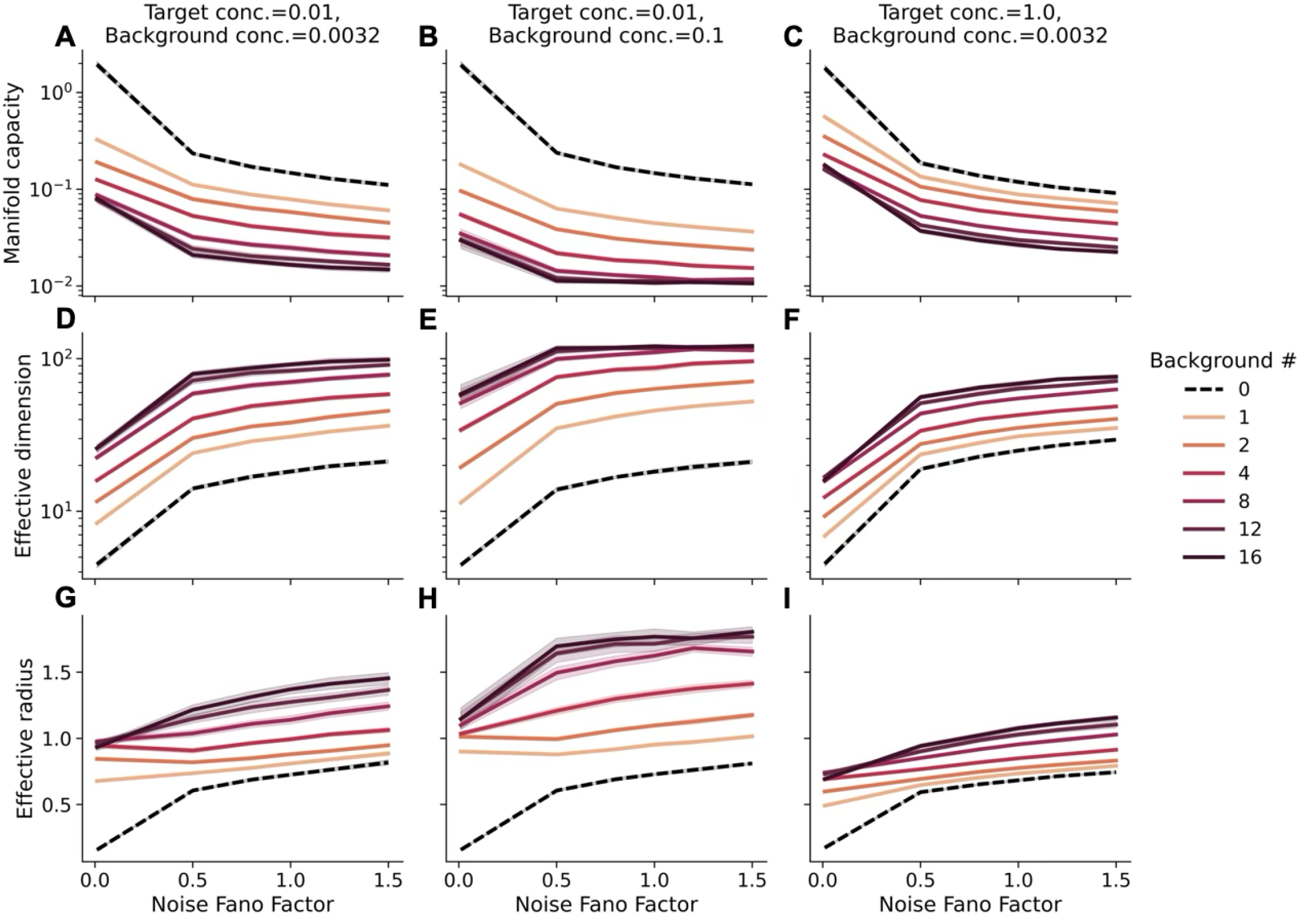
Neural manifold capacity and effective geometry of glomerular representation depend on relative strengths of neural and background noise. (A–C) Manifold capacity, (D-F) effective dimension, and (G-I) effective radius as a function of neural noise strength (controlled by the Fano Factor), background complexity, and cue/background concentrations. Black dashed line represents the effect of only neural noise and target sensitivity. Shaded region represents 95% confidence interval. The theoretical maximum capacity is 4.

The decoder and behavioral analyses showed that lowering target concentration impaired performance broadly across all background conditions, whereas increasing background complexity had comparatively modest effects. Manifold analysis revealed that these manipulations altered representation geometry in fundamentally different ways. Under weak background and strong target conditions, increasing the number of background odors had relatively little effect on manifold geometry: the cue manifolds remained compact and well separated, while effective dimension and radius changed only modestly (Figure 4C,F,I). Thus, weak backgrounds minimally distorted the structure of the cue representations even as target concentration declined. By contrast, lowering target concentration primarily shifted manifold capacity downward (Figure 4A, C), indicating that a weaker signal reduced the separability of the cue manifolds across all background conditions. This geometric trend mirrors the behavioral and linear decoding results, where decreasing target concentration shifted performance downward while maintaining similar slopes across background conditions. When background concentration was high (0.1), however, increasing background complexity substantially expanded effective dimension and radius while reducing manifold capacity (Figure 4B,E,H), indicating that strong backgrounds broadened the cue manifolds and increased overlap between representations far more than weak backgrounds.

We can isolate the effect of target concentration on the geometry by comparing the first and third columns of the figure. We observe the stronger target concentration (Figure 4C,F,I) results in more “compressed” colored curves, meaning the impact of adding more background odors is diminished compared to weak target concentration (Figure 4A,D,G). This can be simply interpreted as a stronger signal results in neural manifolds that remain compact and well separated even as background noise is added.

Next, we can isolate the effect of background concentration on the geometry by comparing the first two columns of the figure. We found that holding target concentration fixed, both weak and strong background concentration produced a similar relationship between the background complexity and effective dimension (Figure 4D,E). Thus, we can interpret the effect of adding background odors at any strength as increasing the number of dimensions along which the odor manifold varies. The magnitude of variability, however, depended on the concentration of background odors, and this is reflected in the increased effective radius of the manifolds at higher background concentration (Figure 4G,H). Together, these analyses provide a geometric explanation for the behavioral and decoding model results. Decreasing target concentration reduces the separability of cue manifolds by increasing both the magnitude and dimensionality of background-induced variations. By contrast, weak background mixtures leave glomerular representations relatively compact and minimally distorted in terms of effective radius, while substantially stronger backgrounds produce the manifold expansion and overlap characteristic of a background-limited regime. Lastly, the addition of background odors results in more dimensions of manifold variability regardless of the background concentration, but a sparse linear readout may pick out a subset of signal-relevant directions and ignore the additional dimensions of variability. Only the change in effective radius remains to influence the separability, and this effect is much stronger with strong background concentration. Thus, the manifold analysis predicts that a background-dominated regime would be mediated by an increased effective manifold radius.

## DISCUSSION

### Odor detectability with decreasing concentration

Our results show that odor discrimination performance declined systematically as target concentration decreased, but remained surprisingly insensitive to background mixture complexity across the tested range of conditions. Although background odors are expected to interfere with perception through overlapping glomerular activation^19,22,42–44^, reducing target concentration did not selectively impair performance in more complex mixtures (Figure 2F). Instead, performance declined similarly across background conditions, indicating that weak target signals became limiting before background interference strongly disrupted discrimination.

This result helps clarify how concentration-dependent limits on odor perception emerge in complex sensory environments. Detection thresholds for monomolecular odors are known to vary widely across many species, including humans^45–47^, presumably depending on the cognate receptors, but how these thresholds behave in complex mixtures has remained largely unexplored. Our findings suggest that, under the conditions tested here, olfactory representations remain relatively robust to increasing background complexity even as target signals weaken. Rather than entering a strongly background-limited regime, behavioral performance was primarily constrained by the detectability of the target odor itself. Consistent with prior work, some background odors produced substantially greater masking of target representations than others when estimated from published glomerular datasets (Fig. S6). However, the degree of masking changed only weakly across target concentrations in the glomerular model, suggesting that the pronounced behavioral dependence on concentration cannot be explained solely by concentration-dependent changes in target-background masking.

### Speed-accuracy tradeoff

The relationship between sampling duration and behavioral accuracy further supports the interpretation that performance was limited by weak sensory signals rather than by background interference. At high target concentrations, mice benefited from longer sampling times, consistent with a classical speed-accuracy tradeoff in olfactory decision-making^31,48,49^. Under these conditions, longer odor sampling likely allowed animals to accumulate additional sensory evidence, improving discrimination performance (Figure 2C-D).

As target concentration decreased, however, the relationship between sampling duration and performance became progressively weaker. Although discrimination accuracy declined substantially at low concentrations, longer sampling no longer improved performance, suggesting that additional sampling provided little useful information in this signal-poor regime. Consistent with this interpretation, sampling times modestly decreased as target concentration declined (Figure 2B), similar to observations in related odor mixture discrimination tasks^30^. Together, these results suggest that when target signals become weak, neural variability limits the quality of sensory evidence itself, such that extending sampling duration cannot substantially improve discrimination accuracy.

### Target odor signal-to-noise limits performance

The weak dependence of behavioral performance on background complexity suggests that, under our task conditions, odor discrimination was limited primarily by target detectability rather than by strong mixture interference. This result is not trivial given the known nonlinearities of olfactory coding. Neural responses throughout the olfactory system vary strongly with odor concentration^34,45,50^, and odor mixtures produce complex receptor-level interactions including suppression and antagonism^33,51,52^. Moreover, concentration-dependent odor identity decoding differs across olfactory bulb output channels, with tufted cells supporting more concentration-invariant decoding than mitral cells^53–56^. These observations, together with prior modeling showing that nonlinear mixture interactions can reduce component decodability^57^ suggest that weak target odors embedded within mixtures could become increasingly vulnerable to masking by background activity.

To understand why this did not occur behaviorally, we analyzed a biophysical model of glomerular activity across different signal and background regimes. Decoder performance best matched behavior in a weak-background regime where target odors activated relatively sparse glomerular representations and background odors produced limited overlap with target-selective glomeruli. Under these conditions, discrimination performance declined primarily because Poisson-like neural variability progressively obscured weak target signals. In contrast, stronger background inputs produced the expected background-limited regime, where increasing mixture complexity substantially impaired decoding as target concentration decreased.

These results suggest that sparse sensory representations help preserve odor discriminability even in complex mixtures. Because individual odorants activate relatively restricted subsets of glomeruli, accurate decoding can be achieved by selectively weighting a limited set of informative inputs while largely ignoring irrelevant background activity. Consistent with this interpretation, sparse readouts better reproduced behavioral performance than dense decoders pooling broadly across glomeruli, and the resulting sparsity levels aligned with experimentally measured glomerular response sparsity^34,38^. This interpretation extends earlier work showing that olfactory bulb representations can support robust linear decoding of odor mixtures^22^ across a broader range of target concentrations and background conditions.

Importantly, our results indicate that early olfactory representations are already sufficiently informative to support robust odor mixture discrimination through simple linear decoding. This interpretation is consistent with our re-analysis of published OSN and piriform cortical bouton recordings^58^, as well as recent simultaneous recordings suggesting greater mixture discriminability in olfactory bulb than piriform cortex representations^59^. However, this does not imply that downstream nonlinear processing is unnecessary in all contexts. Rather, our results suggest that under the conditions studied here, olfactory bulb representations remain sufficiently separable for simple linear readout, whereas stronger interference regimes may benefit from additional cortical transformations^60^. Together, these findings support a model in which downstream cortical circuits may refine, contextualize, or associate odor representations, while olfactory bulb representations remain sufficiently separable to support robust odor discrimination under the conditions examined here.

### Geometry of odor manifolds

To understand how neural variability and background odors alter odor discriminability at the population level, we analyzed the geometry of glomerular activity manifolds generated by the model^41^. This framework provides a geometric description of how easily odor representations can be linearly separated and how different noise sources distort the underlying sensory representation.

Manifold analysis revealed that neural variability and background mixtures affected representation geometry in qualitatively different ways. Poisson-like neural noise consistently reduced manifold separability by increasing the effective dimensionality and spread of odor representations (Figure 4). In contrast, weak background mixtures produced comparatively modest geometric distortions, leaving odor manifolds relatively compact and well separated even as target concentration decreased. Only under stronger background conditions did increasing mixture complexity substantially expand manifold structure and increase overlap between odor representations, consistent with the emergence of a background-limited regime.

This geometric interpretation closely mirrors the behavioral and decoding results. Decreasing target concentration reduced discrimination performance broadly across all background conditions, consistent with neural variability globally degrading separability of the target representations. At the same time, the relatively weak effect of background complexity behaviorally corresponded to the minimal manifold distortions produced by weak background mixtures in the model. Together, these results suggest that sparse olfactory representations preserve robust odor separability across complex mixtures until weak target signals and neural variability become the dominant limitations on discrimination performance.

### Future work

Our results generate several experimentally testable predictions about how sensory variability and background structure shape odor discrimination. First, glomerular population recordings across target concentrations should reproduce the behavioral concentration-performance relationship when analyzed using linear decoders. Second, trial-to-trial decoding variability should be dominated by neural noise aligned with the target representation rather than by large background-dependent distortions of the sensory code. Finally, sufficiently strong background inputs or substantially weaker target concentrations should eventually drive the system into a truly background-limited regime characterized by increased representational overlap and reduced linear separability.

An important next step will be to determine how these principles extend to more naturalistic olfactory environments. Although our mixtures contained up to 16 odor components, natural odor scenes can contain substantially larger numbers of volatile compounds. However, many natural scenes may still possess relatively sparse effective structure, with only a subset of high-concentration odorants dominating sensory representations. Future experiments using stronger backgrounds or hierarchically structured “mixtures of mixtures” could help determine when background interference becomes fully masking and whether the independence between target concentration and background complexity persists under these conditions.

More broadly, simultaneous recordings across olfactory bulb and cortical circuits during odor-guided behavior could clarify how representational geometry evolves across processing stages. In particular, relating manifold structure and decoder performance directly to behavioral variability may help constrain computational models of olfactory perception^41^. An important direction will be identifying the biophysical and circuit-level mechanisms that preserve separability in olfactory representations despite background variability. One possible contributor is heterogeneity in glomerular response amplitudes and tuning widths, which theoretical and experimental studies in sensory cortex suggest can reshape representational geometry by increasing centroid separation or decorrelating manifold structure^61^. Similar mechanisms may contribute to maintaining compact and linearly separable odor representations in the olfactory bulb.

### Concluding remarks

In summary, our experiments reveal that the effects of target concentration and background complexity on odor discrimination are largely independent. Modeling identifies a regime of strong neural noise and weak background interference that reproduces this behavioral robustness, explaining why faint targets remain detectable even in complex mixtures. By combining behavioral data, biophysical modeling, and manifold analysis, we show that glomerular representations of odor mixtures remain sparse and linearly decodable under challenging conditions. These results imply that the olfactory bulb provides a robust substrate for odor identity discrimination, while downstream cortical areas may primarily refine or contextualize this information, a direction for future investigation.

### Limitations of the study

Our study has several important limitations. In the behavioral experiments, mice discriminated between two target odors embedded within artificial background mixtures. Natural odor environments are often more dynamic, containing fluctuating concentrations, temporal structure, and correlations between odor sources that were not represented here. In addition, our task involved only a small number of behaviorally relevant odors. It remains possible that performance would be more strongly affected in environments containing larger numbers of salient odor sources that compete for behavioral relevance.

Our modeling analyses focused on representations at the level of olfactory bulb glomeruli and on the performance of linear readouts. Although the modeled receptor sensitivity distribution is consistent with experimental characterization of olfactory receptors, we do not have a method to calibrate between simulated odor mixture and experimental odor concentrations. Thus, we empirically search for a concentration regime in the model where decoding matches behavior performance qualitatively and retrodictively compare the population response sparsity to glomerular response data.

To capture the 0-background performance drops in the target-dominated regime, we needed neural noise in the response. We assumed Poisson-like neural noise model without correlations across glomeruli, though real glomerular responses are likely to display noise correlations. In terms of readout, downstream regions may transform glomerular representations through nonlinear computations, learning-dependent plasticity, recurrent network dynamics, and statedependent modulation. Such processes could improve the robustness of odor representations or alter how information is extracted from glomerular activity, potentially influencing behavioral performance in ways not captured by our framework.

Our manifold analyses focused on the geometry of static population representations and linear separability. We did not examine how temporal dynamics within sniff cycles, representation learning, noise correlations, or cortical feedback reshape these manifolds. Incorporating these factors may reveal additional trends in the geometric features that contribute to odor discrimination.

Our results support the conclusion that neural noise, rather than background interference, is the primary factor limiting odor discrimination under the conditions examined here. However, under more complex odor scenes, olfactory processing may enter a different regime in which background interference becomes a dominant constraint on performance. The target-dominated regime identified here may therefore represent one portion of a broader continuum of olfactory operating conditions.

## RESOURCE AVAILABILITY

### Lead contact

Requests for further information and resources should be directed to and will be fulfilled by the lead contact, Venkatesh N. Murthy.

### Materials availability

This study did not generate new materials.

### Data and code availability

- This paper analyzes prior data from DOI [https://doi.org/10.7554/eLife.80470]^34^ and [https://doi.org/1020-17124-5]^58^, which are available upon reasonable request to the corresponding authors and [https://doi.org/10.7554/eLife.80470]^38^, which is available under the data availability section.
- Code for glomerular response simulation and decoding has been deposited on GitHub under [https://github.com/george-tog/sparse_inputs] and is publicly available as of the date of publication.
- Code for the manifold geometry analysis will be made publicly available upon publication of Chou et. al 2025^41^.
- Any additional information required to reanalyze the data reported in this paper is available from the lead contact upon request.

## ACKNOWLEDGMENTS

We thank Jacob Zavatone-Veth for vital feedback on an earlier version of the manuscript, and Joseph D. Zak for access to published data we used for analysis leading to Figure S6.

This work has been made possible in part by a gift from the Chan Zuckerberg Initiative Foundation to establish the Kempner Institute for the Study of Natural and Artificial Intelligence at Harvard University. F.P. was supported by the Harvard Center for Brain Science (CBS)-NTT Fellowship Program on the Physics of Intelligence. G.C. was supported by a graduate fellowship from the Kempner Institute. H.M. was supported by an NIH F31 Fellowship (DC023435).

## AUTHOR CONTRIBUTIONS

Conceptualization, V.K. and V.N.M.; investigation, H.M., G.C., C.T., N.H., J.H., F.P. and V.K.; formal analysis, G.C., SY.C. and V.N.M.; writing (original draft), H.M. and G.C.; writing (review and editing), F.P., SY.C, V.K. and V.N.M.; resources, V.N.M.; supervision, V.K. and V.N.M.; and funding acquisition, H.M., G.C., F.P., SY.C. and V.N.M.

## SUPPLEMENTAL INFORMATION INDEX

Document S1. Figures S1-S8

## STAR METHODS

### Key resources table

Included as attachment

### Experimental model and study participant details

Eleven male adult Black6/J (000664) were trained on the behavioral task. All animals were obtained from Jackson Laboratories and maintained within Harvard University’s Biological Research Infrastructure (BRI). Mice were housed in an inverted 12 h light cycle and fed ad libitum. Animals were 2–12 months old at the time of the experiments. All experiments were performed in accordance with the guidelines set by the National Institutes of Health and approved by the Institutional Animal Care and Use Committee at Harvard University.

## Method details

### Behavioral Apparatus

The olfactometer (S1 was custom built in house using a series of clear odor vials, soft tubing, and solenoid valves. The behavior chamber consisted of a custom built sound proofing box, an odor release port utilizing an IR beam break emitter and sensor, and two water releasing ports utilizing capacitive lick sensors and solenoid controlled water release.

Odor port activation, odor release, and monitoring of licking and water rewards were controlled using computer interface hardware (data acquisition boards and I/O controllers, National Instruments) and custom software written in LabView. The mouse was continuously monitored using a web camera during behavior sessions.

### Odors Used

The following odors were used as target odors: methyl tiglate and valeraldehyde. For supplemental analysis (Figure S4) propyl acetate and allyl butyrate were used as targets.

The following odors were used as background odors: pentyl acetate, isoamyl acetate, ethyl tiglate, propyl tiglate, methyl valerate, ethyl propionate, octanol, pentanol, heptanal, isobutyraldehyde, hexanone, heptanone, methyl butyrate, ethyl butyrate, ethyl hexanoate, and hexanal.

### Training Protocol

Water-restricted mice were trained in a freely moving two-alternative forced choice (2AFC) task to discriminate between two target odors mixed within variable odor backgrounds. Each trial began when the animal entered a central odor delivery port, releasing a 1-second odor pulse. Following odor presentation, mice had up to 20 seconds to respond by licking one of two side ports. Correct responses were rewarded with a single drop of water (∼ 2 µL). Once a choice was made, the trial ended, and a new trial began upon reentry to the odor port. Mice were not permitted to switch choices after selecting a port.

Mice were trained in a procedure similar to that described by Rokni et al.^19^. During the initial phase of training, animals learned to discriminate between two monomolecular target odors in the absence of background odors. Once they achieved a performance criterion of ≥75% correct responses, background odors were introduced.

These backgrounds consisted of randomly selected subsets from a pool of 16 monomolecular odorants. Individual background odors were presented at a fixed concentration of 5% (v/v) in all trials. Background mixtures were generated by opening a specified number of odor solenoids connected to separate vials containing the monomolecular background odors, allowing the selected odors to mix within the delivery tubing. This approach maintained a constant concentration for each individual background component regardless of the total number of components in the mixture.

Background complexity was initially drawn from an exponential distribution favoring mixtures with few background components. Over successive training sessions, this distribution was gradually flattened until each level of background complexity, from 0 to 16 components, was equally probable. Under this uniform distribution, mice could encounter up to 65,536 unique background odor combinations.

Mice were considered experts after reaching ≥75% accuracy under uniform background conditions. This typically took anywhere from 2-7 days to achieve after starting the uniform background condition. Following expert performance, mice were further tested with the training concentration for to evaluate the performance at the training concentration without learning effects. Mice were then tested on subsequently lower odor concentrations. Each mouse completed approximately 150–300 trials per day across 3–5 sessions per concentration level. The target odor concentrations were decreased by orders of magnitude across sessions, ranging from 5% to 0.005% (v/v).

For supplementary analysis, instead of sequentially exposing mice to decreasing target odor concentrations, mice were presented with probe trials of lower concentrations interleaved between trials of the training concentration.

### Quantification and statistical analysis

#### Data Analysis

i. **Sample and action time measurement:** Sample time was defined as the total duration during which the odor port beam-break sensor was activated within a trial. This included multiple entries into the odor port, but only the periods when the mouse was inside the port contributed to the sample time. Action time was defined as the interval between the final odor port beam-break and the subsequent activation of either lick port. Periods between earlier sampling bouts were not included in the action time.
ii. **Trial filtering:** For analyses investigating how performance changed across increasing background difficulty and learning (Figure 1), two mice were excluded because their training schedules differed from the standardized regime used for the remaining animals. Mice included in this analysis completed at least six days of trials on the most difficult background complexity at the training target odor concentration, and data from the first three training days were retained to capture within-animal learning. In contrast, for analyses examining steady-state performance across concentrations and background complexities (Figure 2), the first three days of recordings at the “training” concentration were excluded. These sessions represented initial learning, whereas all subsequent sessions were considered “test” days and reflected expert-level performance.

### Behavioral Fitting and Statistical Analysis

For analyses of performance across training and background complexity (Figure 1C), a two-way ANOVA was used to test main effects of day and background complexity, as well as their interaction.

To assess whether behavioral timing metrics (sample and action time) changed across days (Figures 1D, S2C) or target odor concentrations (Figures 2B S2D), we compared trial-wise distributions using two-way ANOVAs (for day × background complexity) and log-linear regressions (for concentration). Log-linear regressions were of the form y = β_0_ + β_1_ log_10_(x), where y was the behavioral measure and x the target concentration; slope coefficients (β_1_) were tested for significance using t-tests.

Behavioral performance across target odor concentrations (Figure 2A) was summarized using a logistic (sigmoid) function:

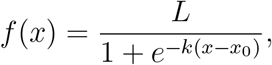

where L denotes the asymptotic maximum performance, k the slope (steepness) of the function, and x_0_ the inflection point corresponding to the concentration at which performance reaches half of L. Logistic functions were fit to aggregated performance data using nonlinear least-squares regression and are shown for descriptive visualization. Goodness of fit was quantified using the coefficient of determination (R^2^).

To statistically assess the effect of odor concentration on behavioral performance while accounting for repeated measurements within animals, we fit a trial-level mixed-effects logistic regression model. Trial outcome (correct or incorrect) was modeled as a binomial response with log_10_(concentration) as a fixed effect and animal identity included as a random intercept. Statistical significance of the concentration effect was assessed using Wald z-tests on the fixed-effect coefficient.

To quantify the concentration-dependence of the speed-accuracy relationship (Figure 2C-D), we estimated the slope of performance as a function of odor sampling time, separately at each of the six target odor concentrations. Trials were combined across animals into bins of sampling time (bin width 25 ms for durations up to 275 ms, wider 50 ms bins at longer durations), and the mean performance and trial count were computed for each bin. Bins with zero trials were excluded. For each concentration, a binomial generalized linear model with a logit function was fitted using the binned trial counts as the response:

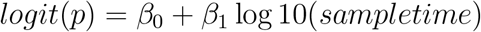

where p is the probability of a correct response and sample time is the bin center in milliseconds. This approach treats each bin as a binomial observation (N correct out of total trials), weighing bins by trial count.

The slope β_1_ was was converted to units of percentage points per log_10_ (ms) by evaluating the fitted model at sample times of 75 ms and 300 ms (a range spanning the central portion of the data). Uncertainty on this derived slope was estimated by parametric bootstrap: 10,000 draws of (β_0_, β_1_), and the mean and SD of this distribution were reported as the slope estimate and SD respectively. Statistical significance was assessed from the Wald z-test on β_1_

For analysis of performance across background concentrations (Figure 2F), background odor numbers (0–16) were grouped into eight equal-width bins (0–2, 2–4, 4–6, 6–8, 8–10, 10–12, 12–14, 14–16) to reduce variability and visualize overall trends across background complexities. Slopes and R^2^ values for each concentration were derived from least-squares fits across per-animal means (e.g., slopes of −0.002 to −0.004 with corresponding R^2^ values of 0.01–0.04).

Differences in overall performance across odor concentrations (Figure 2F) were tested using a one-way ANOVA followed by Tukey’s Honest Significant Difference (HSD) post-hoc tests. This analysis evaluated whether mean performance differed between consecutive concentration levels (e.g., 5.0% vs. 2.5%, 2.5% vs. 0.5%, etc.).

For visualization, shaded regions represent ±SEM across animals unless otherwise specified. All statistical analyses were performed in Python.

### Biophysical Model

We simulate glomerular responses using the biophysical model from^33^. We summarize the model here and refer the interested reader to the original work for justification of the modeling choices.

A mixture of K monomolecular odorants *X*_1_, *X*_2_, … , *X*_*K*_ with concentrations *C*_1_, *C*_2_, … , *C*_*K*_ is presented to an array of odor receptor neurons. The odorants competitively bind to odor receptors, which are parameterized by two vectors: binding affinity 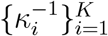 and activation efficiency 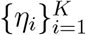. The parameters are sampled i.i.d. (per odor indexed by i) from a bivariate log-normal distribution:

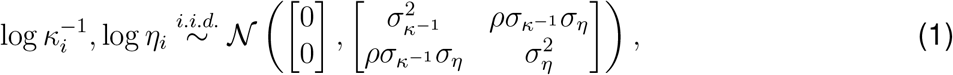

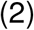

where the correlation *ρ* between log-sensitivity and log-activation efficiencies is a tunable parameter:

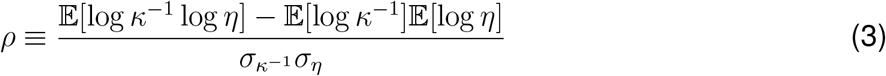

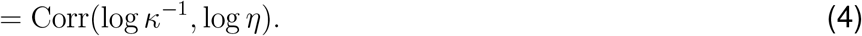

When *ρ* = 1, the rank order of sensitivity will perfectly match the rank order of activation efficiency since logarithm is a monotone function. When the correlation is less than 1, it is possible for odorant *A*, which activates receptor *i* less efficiently than odorant *B*, to bind with higher affinity than odorant B. This effect is termed “antagonism” and can be considered a normalization of the population response that improves detection of individual components^33,34^. To match experimentally measured sensitivity ranges, we set 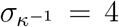to obtain six orders of magnitude separation between the most and least sensitive receptor types.

The ORN response to a mixture of K odorants with total concentration 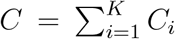 and fractional concentrations *β*_*i*_ = *C*_*i*_/*C* is given by:

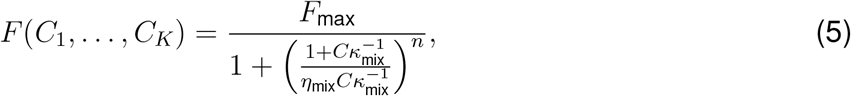

where the effective mixture parameters are defined as:

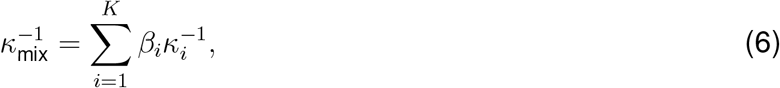

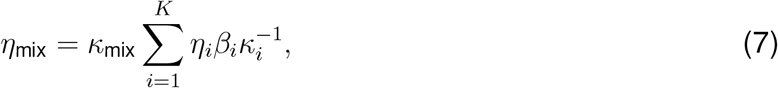

and the Hill coefficient is set to *n* = 4. The maximal response is *F*_max_ = 10, and the mixture response has the form of a monomolecular odor response with effective sensitivity and activation efficiency parameters. We show example glomerular responses to odor mixtures of different background complexities containing each target odor in Figure 3A.

### Decoding Analysis

To approximate the experimental training paradigm, we independently generate 1000 train and test mixtures that contain one of two cue odors and a random selection of up to 16 background odors. The cue odors and the number of background odors in each mixture were sampled uniformly. We generate the glomerular responses from N = 400 glomeruli for each of these mixtures, and add neural response noise 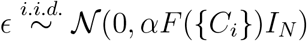. The variance is mean-dependent up to the Fano factor α ∈ [0,∞ ). This parameterization is a continuous approximation of Poisson noise for α = 1, and α serves as a knob to tune the neural noise strength to be sub- or super-Poissonian. The approximation is justified by the fact that Poisson spike counts approach a Gaussian distribution for a large rate parameter, and a glomerular response comprises activity from many converging OSNs over a time window.

We use a logistic regression classifier from the `scikit-learn` package with L1 regularization strength *C* = 1 to fit the training set to binary labels specified by the cue odor present in each mixture. We additionally consider L2 regularization with *C* = 1. The classifier then predicts the cue odor present in the mixtures that elicited each test response, and the fraction of correct predictions is reported as “decoder accuracy” (Figure 3). We use 10 replicates of independently sampled odor mixtures and responses while holding the sampled receptor parameters *κ*^−1^ and *η* fixed across replicates.

In our analyses, we sweep across the following parameters: neural noise Fano factor α ∈ {0, 0.01, 0.5, 0.8, 1.0, 1.2, 1.5}, cue concentration (train/test) *C*_1_, *C*_2_ ∈ {10^−7^, 10^−6.5^, … , 10^−3^}, background concentration *C*_3_, … , *C*_*K*_ ∈ {10^−7^, 10^−6.5^, … , 10^−3^}, and antagonism parameter ρ ∈ {0.5, 1.0}.

### Comparing Behavioral and Decoder Performance

To compare decoder performance in the model to behavioral performance, we summarized the effect of background odor complexity using a ratio-based metric. For each condition, we computed the ratio between performance at the highest and lowest background complexity. Behavioral performance was defined as mean animal success rate, while model performance was defined as decoder accuracy. This metric provides a scale-independent measure of robustness to increasing background odor complexity.

Because the concentration sets used in the behavioral experiments and in the model simulations differed, direct one-to-one concentration matching was not possible. Instead, we compared behavior and models using concentration-rank matching: behavioral and model conditions were independently ordered from highest to lowest cue concentration, and corresponding ranks were matched until one set was exhausted. This approach preserves the relative ordering of concentrations while avoiding assumptions about equivalence between absolute concentration values across experiments and simulations. In addition, in the model simulations many of the highest concentration conditions yielded indistinguishable (flat) decoder performance curves, making them redundant for quantitative comparison; rank matching naturally accommodates this redundancy without over-weighting these conditions.

Agreement between behavioral and model robustness ratios was quantified using the mean absolute error (MAE) and root mean squared error (RMSE) across matched concentration ranks. For each matched rank *i*, we computed the difference between the behavioral robustness ratio 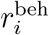 and the corresponding model robustness ratio 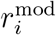. MAE was defined as the mean of the absolute differences across all matched ranks, while RMSE was defined as the square root of the mean squared differences. Lower MAE and RMSE values indicate closer agreement between model predictions and behavioral performance. Differences in model–behavior agreement were assessed using a paired Wilcoxon signed-rank test on absolute errors between behavioral and model robustness ratios across matched concentration ranks.

### Sparsity Analysis

To calibrate our model’s population response sparsity, we compared to glomerular responses (Δ*F*/F ) in two studies. First, 228 glomeruli were imaged during passive exposure to a panel of 32 monomolecular odorants in a single mouse from a previous study, with a range of concentrations from 0.08% to 80% v/v before 16-fold dilution^34^. Second, 1290 glomeruli from eight olfactory bulbs in anesthetized mice were imaged with a 185 odorant panel^38^, and we use the study’s computed sparsity values directly.

To quantify sparsity, we use the complement of the Treves-Rolls sparsity measure:

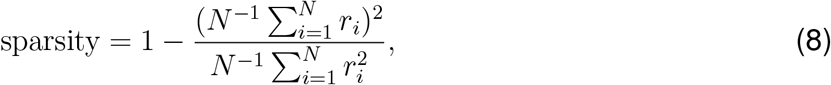

where *N* is the number of neurons and *r*_*i*_ is the firing rate of the *i*th neuron^62^. We take the complement so that a value of 1 − 1/*N* is the sparse limit of a single neuron responding and a value of 0 is the dense limit of all neurons responding uniformly. Because the measure requires non-negative responses, we rectify the Δ*F*/*F* signal in the data. We directly compute the sparsity measure of the biophysical model responses with 228 glomeruli to 185 odorants using the “strong”, “weak”, and “very weak” concentrations of 10^−1^, 10^−2.5^, 10^−4.5^ respectively. For “mixture”, we compute the sparsity using a range of odorant concentrations that uniformly tiles the log-space between (10^−3.5^, 10^−0.5^).

### Glomerular Masking Analysis

To estimate the degree to which background odors interfered with target odor representations at the glomerular level, we computed the masking index *M*(*m,T* ) following Rokni et al. (2014)^19^. Masking quantifies the extent to which a background odor mixture activates glomeruli that are also activated by the target, relative to the target’s own activation at those glomeruli. For each target-activated glomerulus i:

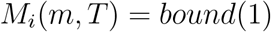

where *a*_*i*_(*T* ) and *a*_*i*_(*m*) are the responses of glomerulus i to the target odor alone and the background mixture alone, respectively. The overall masking value was taken as the mean across all target-activated glomeruli:

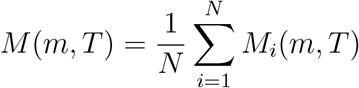

### Experimental data masking

For figures Figure S6B-C, glomerular response profiles were taken from Zak et al. (2020), which imaged 228 glomeruli in response to 32 monomolecular odors in a single animal. Responses were peak Δ*F*/*F* values averaged over 20 frames following odor onset. Negative Δ*F*/*F* values were rectified to zero prior to analysis. Target activated glomeruli were defined as those with Δ*F*/*F* > 0.05 Of the 16 background odors used in the behavioral experiment, 7 were present in the Zak et al. (2020) panel (ethyl tiglate, ethyl propionate, pentyl acetate, methyl valerate, 2-hexanone, methyl butyrate, 2-heptanone) and were used for masking estimation. Background mixture responses were modeled as the linear sum of individual background responses. Masking was computed for all possible combinations of n backgrounds (random sampling of 1000 combinations) and the mean and SD across combinations are reported.

### Model-based masking

To examine how masking varied with target concentration Figure S6D we used the biophysical glomerular model described above. Single-odor response vectors were simulated for target odors and all 16 background odors at target concentrations ranging from 10^−5^ to 10^−2^ and a fixed background concentration of 10^−2^. Masking was computed as above across 1000 random background combinations per complexity level, averaged across 50 independent receptor parameter replicates. The activation threshold was set to the 90th percentile of the target response distribution.

### Sparse, Expansive Cortical Code Analysis

Following^63^, we simulate a sparse, expansive code 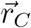 that takes the glomerular model responses 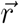 as input. We sample a Gaussian weight matrix 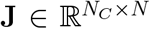 where 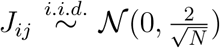 with N glomeruli. The cortical response was modeled as a thresholded linear transformation 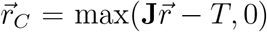, where the threshold *T* was adjusted to achieve the desired fraction of active neurons. In our simulations, we generate responses from 400 glomeruli to odors at 10^−2.5^ concentration to an antagonism parameter *ρ* = 0.5 and uniformly drawn background complexities. We compute 5-fold stratified cross-validation accuracy of an L1-penalized logistic regressor using default hyperparameters on an one vs rest classification with 943 odor mixtures of 1-16 odors. Standard error bars are computed over 16 classification tasks.

### Olfactory Sensory Neuron and Cortical Feedback Decodability Analysis

We analyzed previously published, z-scored calcium imaging responses (Δ*F*/*F* ) of awake mice inhaling mixtures of a standardized odorant panel of 16 monomolecular odorants^58^. In total, responses to 100 mixtures comprising either 1, 4, 8, or 12 odorants were measured in olfactory sensory neurons and cortical feedback boutons. For a fair comparison between areas, we randomly subsampled 120 neurons from each area before fitting classifiers to identify the presence or absence of each the 16 odors. We compute the accuracy of L1-regularized logistic regression classifiers using 5-fold stratified cross-validation without further data preprocessing and average the resulting accuracies across the 16 classification tasks. In Figure S7(A), we plot the mean and standard error of the decoding accuracy for each area.

### Manifold Geometry Definitions

We provide definitions for the effective manifold radius *R*_*M*_ and effective manifold dimensionality *D*_*M*_ in this section following^41^. Let **y** ∈ {−1, +1}^*P*^ be the vector of labels for P neural manifolds. Let the convex hulls of these neural manifolds be denoted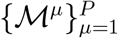 . For a given separating hyperplane with normal vector *w* in the neural activity subspace ℝ^*N*^ , where *N* denotes the number of neurons, the anchor points with respect to the µth neural manifold are defined as s^*µ*^(y, w) = *y*^*µ*^ · x^*µ*^(y, w), where λ, {x^*µ*^(y, w)} are the set of maximizers of

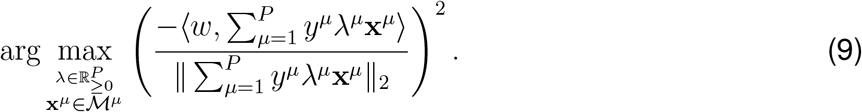

In geometric terms, the anchor point is a representative point on the convex hull of a manifold with respect to a hyperplane. By representative, we mean the closest point with signed distance to the hyperplane because correctly classifying the closest signed point on the convex hull guarantees correct classification of the entire convex hull.

Now, we will consider a random multivariate Gaussian hyperplane normal vector t ∼ *N* (0, I), which induces a distribution of anchor points **s**^*µ*^(**y, t**) that has randomness due to both the labels **y** and the separating hyperplane defined by t. The multivariate Gaussian vector is used to measure the “size” of the convex cone of valid separating hyperplanes, using a measure called statistical dimension^41^. Intuitively, if we draw a random Gaussian vector, the fraction of the time it lies inside or on a convex cone is greater if the cone is larger in angle. The statistical dimension formalizes the size by taking the average squared magnitude of the projection of a multivariate Gaussian vector onto the cone being measured. To connect statistical dimension to capacity, we consider the cone of valid separating hyperplanes. If a representation has high capacity, then manifolds are more linearly separable, and there is a larger cone of valid separating hyperplanes. This would be quantified by a larger statistical dimension. Importantly, this measure does not assume Gaussian neural manifolds.

Now, we introduce matrix notation. Let **S**(**y, t**) ∈ ℝ^*P* ×*N*^ be the matrix of anchor points with **s**^*µ*^(**y, t**) in the µth row. Let the Gram matrix (matrix of inner products of all anchor points) be denoted **G**(**y, t**) = **S**(**y, t**)**S**(**y, t**)^T^. In general, **S**(**y, t**)^T^**G**(**y, t**)^†^**S**(**y, t**) is an orthogonal projection operator onto the subspace spanned by the rows of **S**(**y, t**). The analytic formula for capacity depends on the following field:

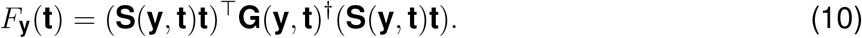

The expectation of this field is the critical dimensionality below which a random projection of the neural manifolds has low probability of being linearly separable.

Next, we will decompose each anchor point into a center and axial component. Define the center component to be the expectation with respect to the anchor point distribution

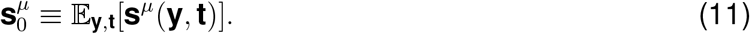

Then, the axial component is defined as

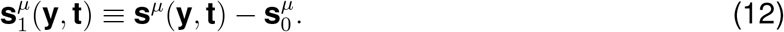

Similar to before, arrange the anchor point components into matrices **S**_0_, **S**_1_(**y, t**) ∈ ℝ^*P* ×*N*^ and denote the component Gram matrices **G**_0_ = **S**_0_**S**^T^_0_ and **G**_1_(**y, t**) = **S**_1_(**y, t**)**S**_1_(**y, t**)^T^. We define the auxiliary fields

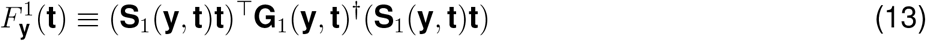

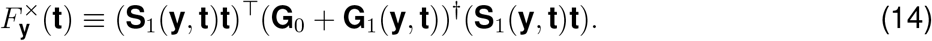

The field 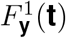 corresponds to the squared norm of the projection of **t** onto the subspace spanned by the axial components of the anchor points.

Finally, we define the effective geometric measures:

1. Effective manifold radius

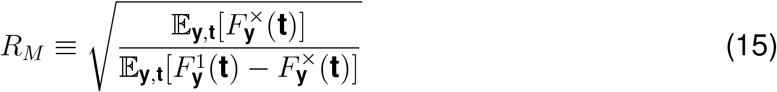
2. Effective manifold dimension

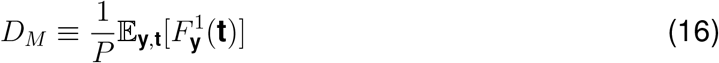
3. Effective manifold utility

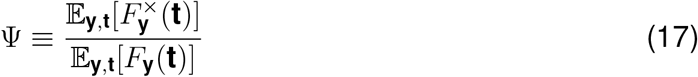

The geometric formula for manifold capacity from the GLUE approach is then

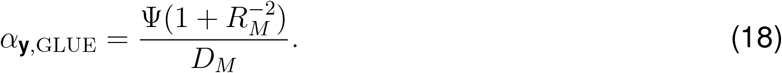

### Capacity Analysis

To better characterize how neural response structure affects odor discriminability beyond simple linear decoding accuracy, we applied the manifold capacity framework. We will first describe the procedure we use to estimate the manifold capacity before providing a concise explanation of the data-driven manifold capacity theory.

We independently generate 10,000 odor mixtures that contain one of two cue odors and a random selection of up to 16 background odors. This time, we allow the backgrounds to be drawn from up to 24 odors to mitigate the effect of only having a single mixture with 16 background odors and 16 mixtures with 15 background odors. We generate glomerular responses for 400 glomeruli under the same biophysical parameter distributions as the decoding analyses; we additionally sample 10 replicates of the biophysical parameters to quantify the uncertainty in our analysis. We consider the set of all neural responses to mixtures containing a given cue odor to be a neural manifold associated with that cue odor. We arbitrarily label one manifold +1 and the other −1. Then, using the “Geometry Linked to Untangling Efficiency” approach^41^, we estimate the manifold capacity, effective manifold radius, and effective manifold dimensionality. In essence, the effective manifold radius (*R*_*M*_ ) describes the extent of variability within each neural response manifold, with smaller radii indicating tighter clustering (less variability). The effective manifold dimensionality (*D*_*M*_) quantifies how many distinct directions of neural response variability are relevant for discriminating between odor categories, with lower dimensionality representations resulting in simpler discrimination. We schematically depict these measures in Figure S8. We sweep across the same glomerulus response parameter sets as in the decoding analyses. We focus our analyses on trends over varying background complexity and neural noise strength at different background and cue concentrations.

Now, we provide a concise explanation of the data-driven manifold capacity. The capacity of the glomerular representation depends on the probability of linear separability after uniform, random projection of the neural response vector to lower dimensions. Formally, let the convex hulls of neural response manifolds be denoted *M*^*µ*^ ⊆ ℝ^*N*^ , with manifold indices *µ* = 1, 2, … , *P* for *P* manifolds (in our case, *P* = 2). Let the associated labels be *y*^*µ*^ ∈ {−1,+1} . The linear classification condition with a hyperplane defined by normal vector *w* ∈ ℝ*N* is *y*^*µ*^ ⟨w, x^*µ*^⟩ ≥ 0 for all µ 1, … , P , x^*µ*^ *µ* ∈ {1,…,*P*},*x*^*µ*^ ∈*M*^*µ*^ .Let Π_*n*_ : ℝ ^*N*^ →ℝ^*n*^ be a uniform, random projection operator, so we can define the following probability of linear separability after random projection to *n* dimensions:

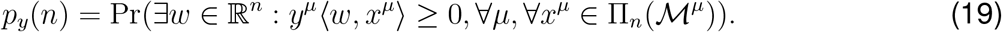

Then, the capacity is defined as

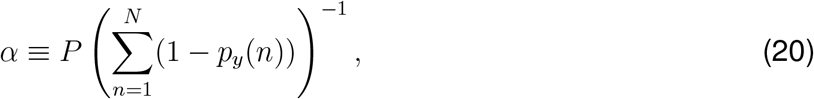

where the summation can be interpreted as the expectation of a random variable with cumulative distribution function *p*_*y*_(*n*). Intuitively, the capacity is the ratio of the number of manifolds to an expected dimensionality below which a random projection of the neural manifolds will no longer be linearly separable. A neural representation that is more suitable for linear classification will have higher capacity.

Directly computing the probability of linear separability is computationally costly, so we use the following analytical formula that both matches simulations and links capacity to geometric properties of neural manifolds. This connection allows us to interpret capacity in terms of the structure in neural population activities.

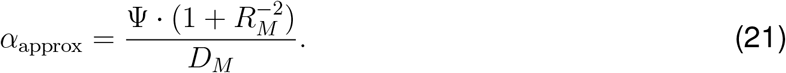

where *R*_*M*_ and *D*_*M*_ are defined in equations 15 and 16 respectively. This analytical formula captures the geometric structure of manifolds through the anchor point distribution, which can be intuitively understood as the representative point on the convex hull of each manifold that is closest to the separating hyperplane. The anchor distribution can be summarized by geometric statistics such as effective manifold radius *R*_*M*_ and effective manifold dimension *D*_*M*_ . Conceptually, the effective manifold dimension measures the number of directions that are spanned by task-relevant intra-manifold variability. The effective geometric statistics are defined so that the case of L2 sphere manifolds with dimensionality d and radius r have geometric statistics R_*M*_ = r and *D*_*M*_=d. The utility Ψ links the effective geometric measures to α_*approx*_ by accounting for how manifolds are organized in the neural representation space. We refer the interested reader to^41^ for further exposition on the data-driven capacity theory and the precise definition of the effective geometric measures.

---

**Figure S1.**
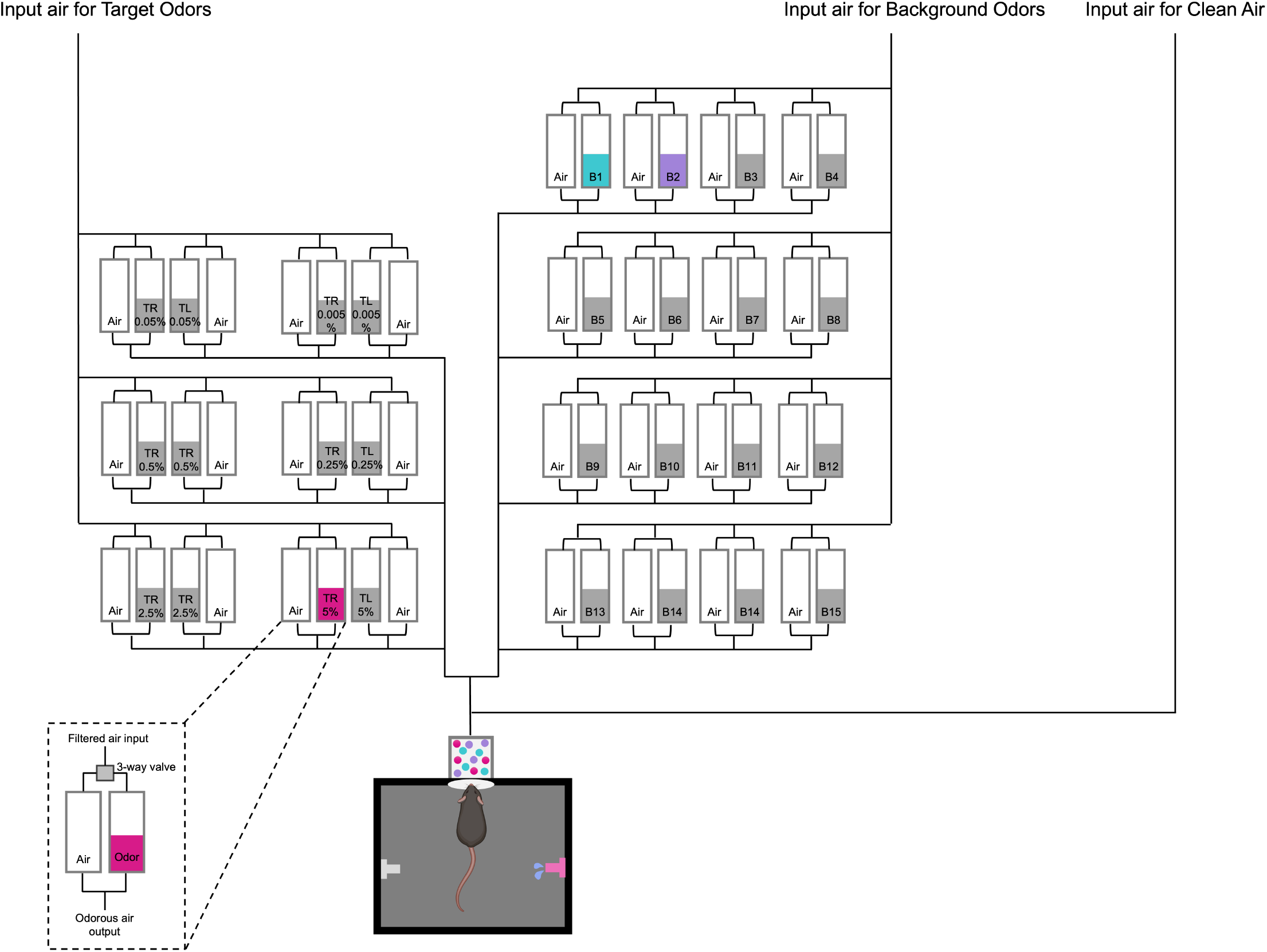
Olfactometer for behavioral experiments An olfactometer was custom built to control precise delivery of odor mixtures to the mouse. This design was based on previous work from^19^. Two main modules controlled the odor release. The first controlled the background odors, allowing for each of the 16 background odorants to be present or absent in any mixture. To keep the total flow constant, each background odor had a matched, clean air control. Each odor-air module had a 3-way valve that either diverted the air through a glass tube containing the background odor or a glass tube containing air. Both paths converged to form the output flow. Odorant mixtures were generated by controlling the 16 3-way valves allowing 2^16^ background odor mixtures. The second main module controlled the target odors. The overall logic, using 3-way valves to divert input air to either a clean air or odorous air tube, remained the same. Only two target odors, matched in concentration, were used in a given session. Target odors were mutually exclusive. Together the odorous air for the target and background odors merged before traveling through a 2-foot long tube of 1/8 inch diameter towards a final 3-way valve. This valve toggled between clean air, and the odorous air containing the target and background mixtures. Upon interruption of the infrared (IR) beam at the odor delivery port by the mouse, the 3-way valve directed odorous air to the port for 1 second. Following odor presentation and until the onset of the subsequent trial, clean air was delivered.

**Figure S2.**
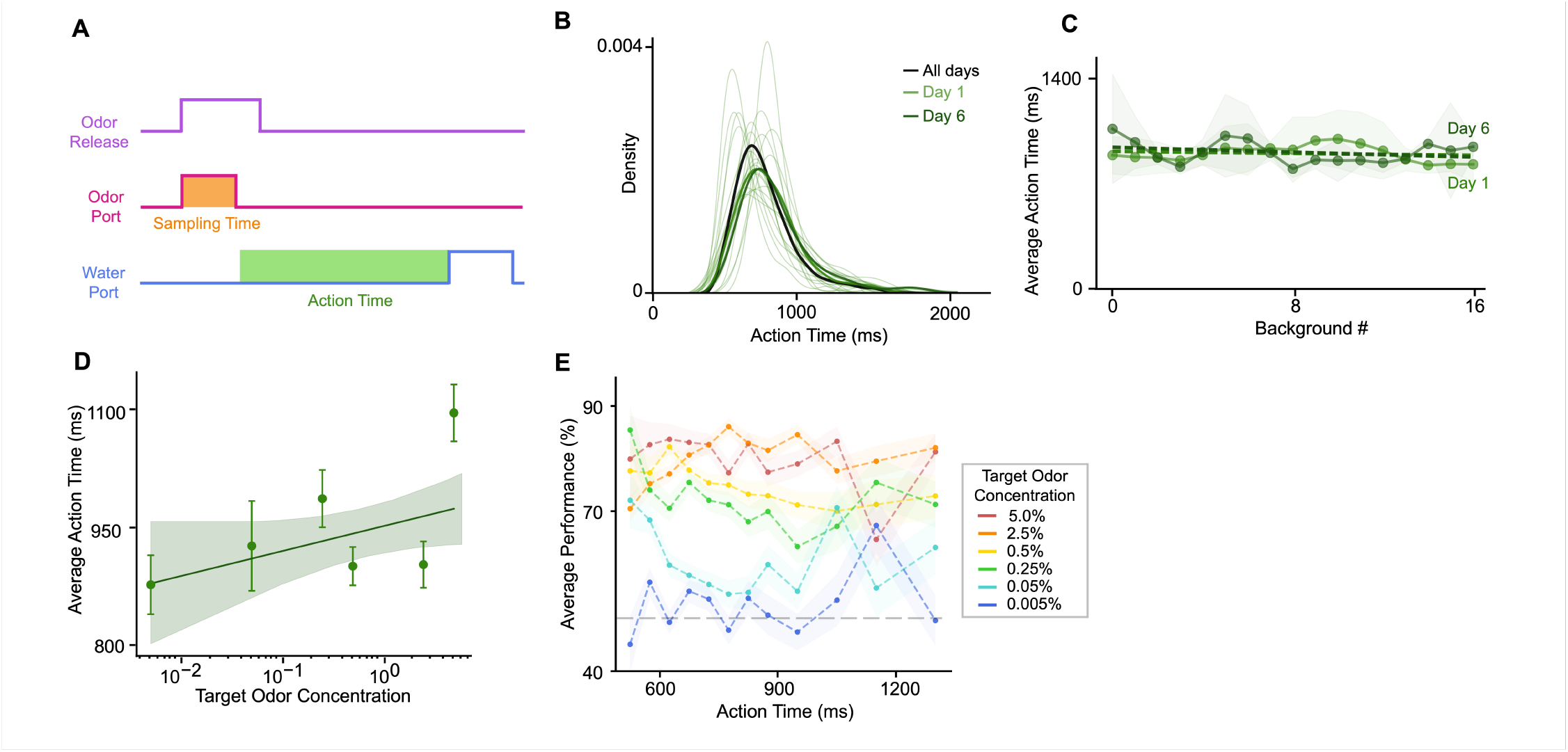
Action Time does not correlate with performance. (A) Schematic representation of the trial structure, highlighting sample and action time (B) Trial-wise distributions of action times at the highest background complexity. Thin lines show individual-day distributions, the bold black curve shows the overall average, and light versus dark curves indicate Day 1 and Day 6 of training. (C) Action time remained stable across days (Day 1 = 890.3 ms, Day 6 =902.8 ms) (two-way ANOVA for day × background complexity, p = 0.69). (D) Action time trended upward with concentration but was not statistically significant (log-linear regression, p = 0.087).(E) Across target odor concentrations, performance did not vary systematically with action time (two-way ANOVA, p = 0.65).

**Figure S3.**
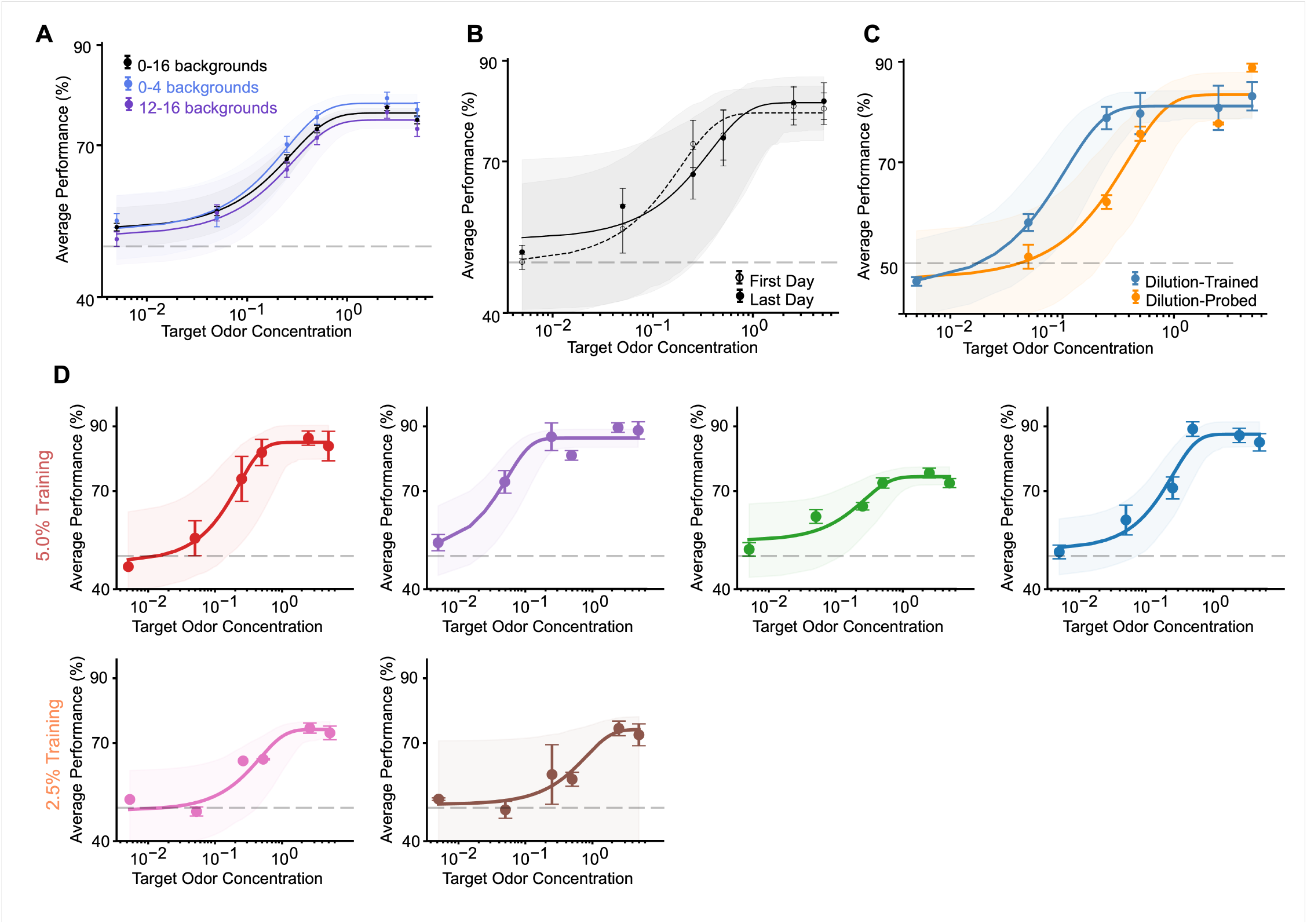
Task performance across different background complexity, training, and probe conditions. (**A**) Psychometric curves across background complexities. Low background conditions (0–4) showed slightly higher asymptotic performance and a steeper logistic slope than high backgrounds (12–16), indicating better discrimination under low-complexity conditions. (**B**) Psychometric curves on the first and last day of training. Logistic fits show an increased maximum performance and a modest change in threshold concentration, consistent with improved discrimination accuracy over training. (**C**) Psychometric curves for probe and control conditions. The probe condition showed a shallower slope and a higher threshold concentration than the control, indicating reduced sensitivity under probe conditions. (**D**) Individual-animal psychometric functions showing consistent maximal performance and similar concentration sensitivity across mice. All panels display logistic best-fit curves with shaded 95% confidence intervals.

**Figure S4.**
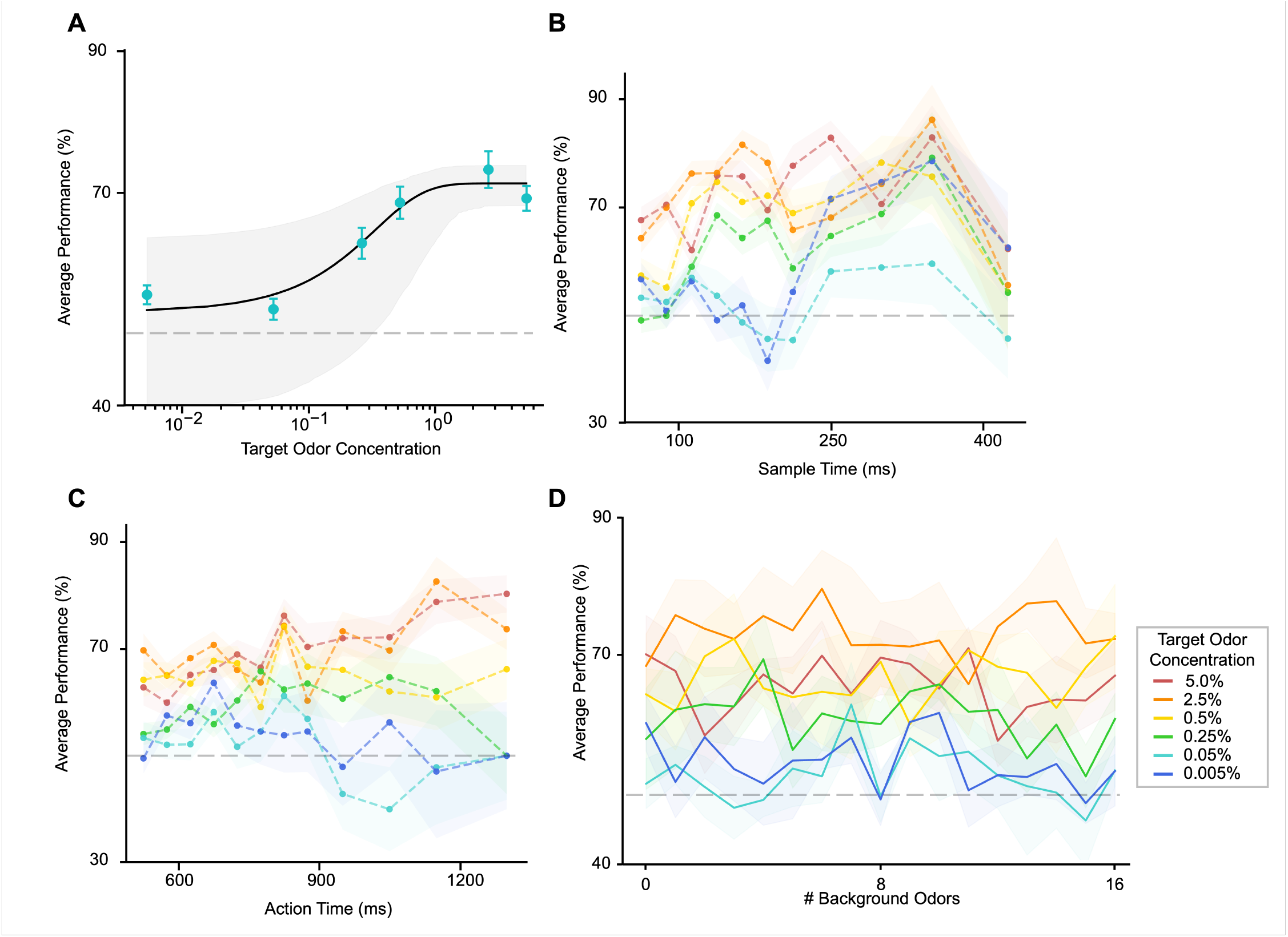
Behavioral Metrics for Alternative Target Odor Pair. Behavioral performance and timing as a function of target odor concentration for alternative target odors, propyl acetate and allyl butyrate, (n = 5). (A) Psychometric curve demonstrating performance as a function of target odor concentration for hard odors. Performance decreased with decreasing concentration, following a sigmoidal trend. Each point represents the mean performance of a single mouse on a single training day. The solid curve shows a logistic fit to the aggregated data (R^2^ = 0.40), with shaded area indicating the 95% confidence interval of the fit. The dashed line denotes chance performance (50%).Performance decreased significantly with concentration (trial-level mixedeffects logistic regression with animal as a random effect, p < 0.001). (B) Sample time showed no effect with increasing target odor concentration (log-linear regression). (C) Action time trended upward with concentration (log-linear regression). (D) Average performance across background complexities at six concentrations of target odor, spanning three orders of magnitude. Gray dashed line represents chance performance (50%). Shaded regions represent ±SEM across animals.

**Figure S5.**
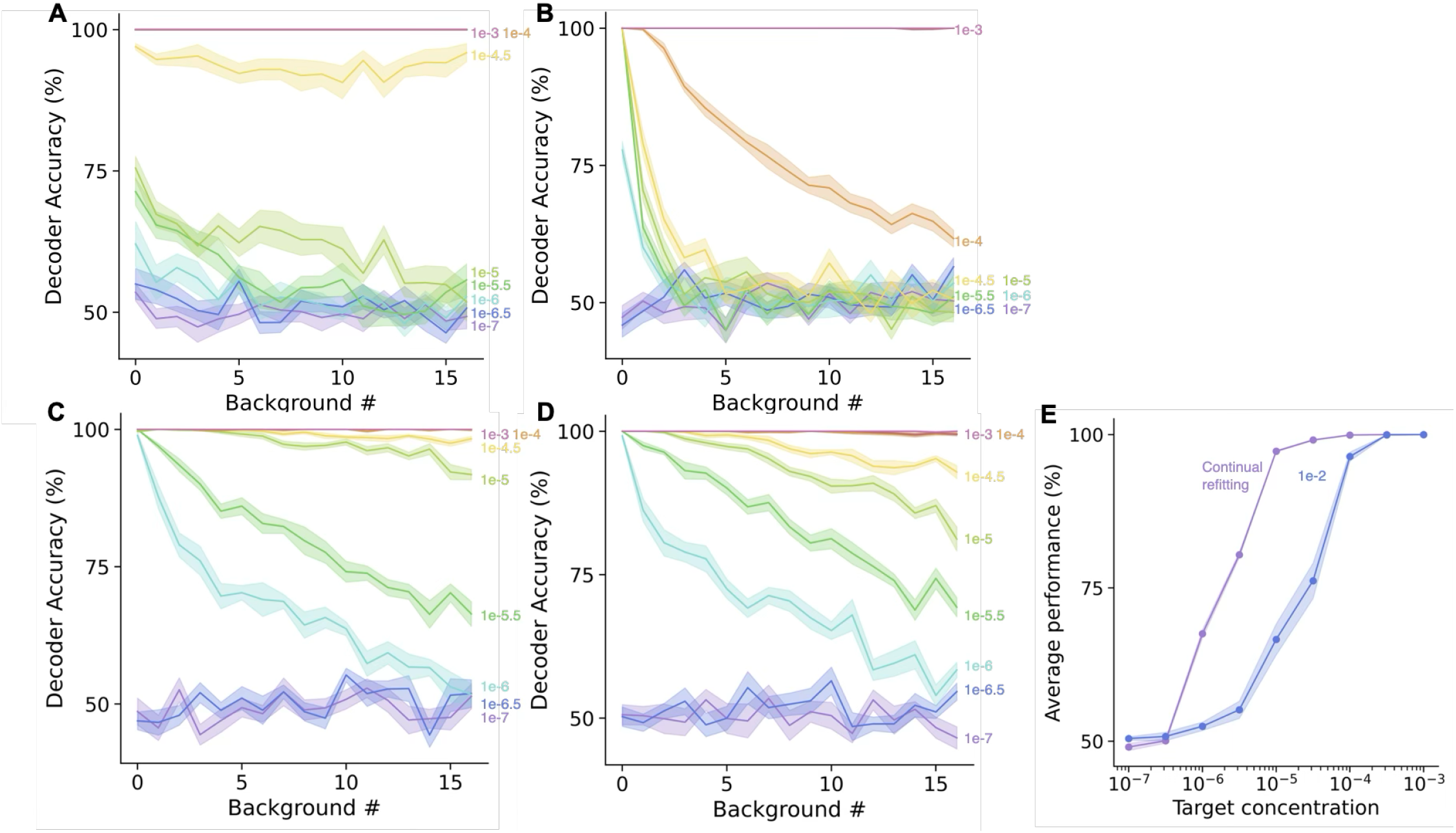
Additional linear decoding settings. First row: Decoding accuracy with (A) L1 regularization, *ρ* = 1.0 (no antagonism), training target concentration 10^−3^, background concentration 10^−3^ (B) L2 regularization, *ρ* = 0.5, training target concentration 10^−2^, background concentration 10^−2.5^. Second row: We use antagonism (ρ = 0.5), L1 regularization, and background concentration 10^−2.5^. (C) Decoders are fit and tested at matched target odor concentrations. This approximates continual learning at each lowered target concentration. (D) The training set consists of 1000 odor mixtures with each target odor concentration for a total of 9000 mixtures. (E) Average performance over all background complexities of a decoder fit and tested at the same target concentration (“continual refitting”) or fit with target concentration 10^−2^ and tested without re-fitting at a range of target concentrations.

**Figure S6.**
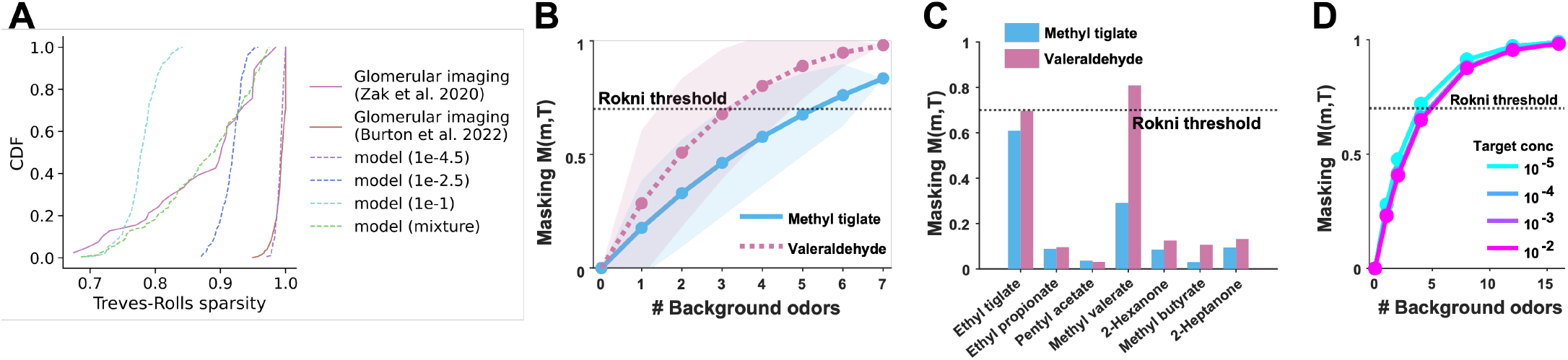
Population response sparsity and masking from experimental data and model. (A) In the data from Zak et al. 2020 ^34^, Treves-Rolls sparsity was computed from the rectified (non-negative) monomolecular evoked response Δ*F*/*F* . See^38^ for how data were pre-processed before sparsity calculation. In the glomerular model, we compute the sparsity using the responses to 185 odors at different concentrations indicated in the legend. The concentrations 10^−2.5^ and 10^−1^ correspond to “weak” and “strong” background respectively. The mixture curve contains odor concentrations uniformly sampled in log-space from (10^−3.5^, 10^−0.5^). (B) Masking index M(m,T) as a function of background complexity for methyl tiglate and valeraldehyde, computed from Zak et al. (2020) glomerular imaging data. Points show mean ± SD across background combinations. Dashed line indicates the Rokni et al. (2014) masking threshold (*M* = 0.7). (C) Single-background masking for each background odor available in the Zak et al. (2020)^34^ dataset. Dashed line indicates the Rokni et al. (2014)^19^ masking threshold (*M* = 0.7). (D) Masking index from the biophysical model across background complexity levels at four target concentrations (10^−5^ to 10^−2^). Background concentration was fixed at 10^−2^. Lines show mean across background combinations and receptor parameter replicates.

**Figure S7.**
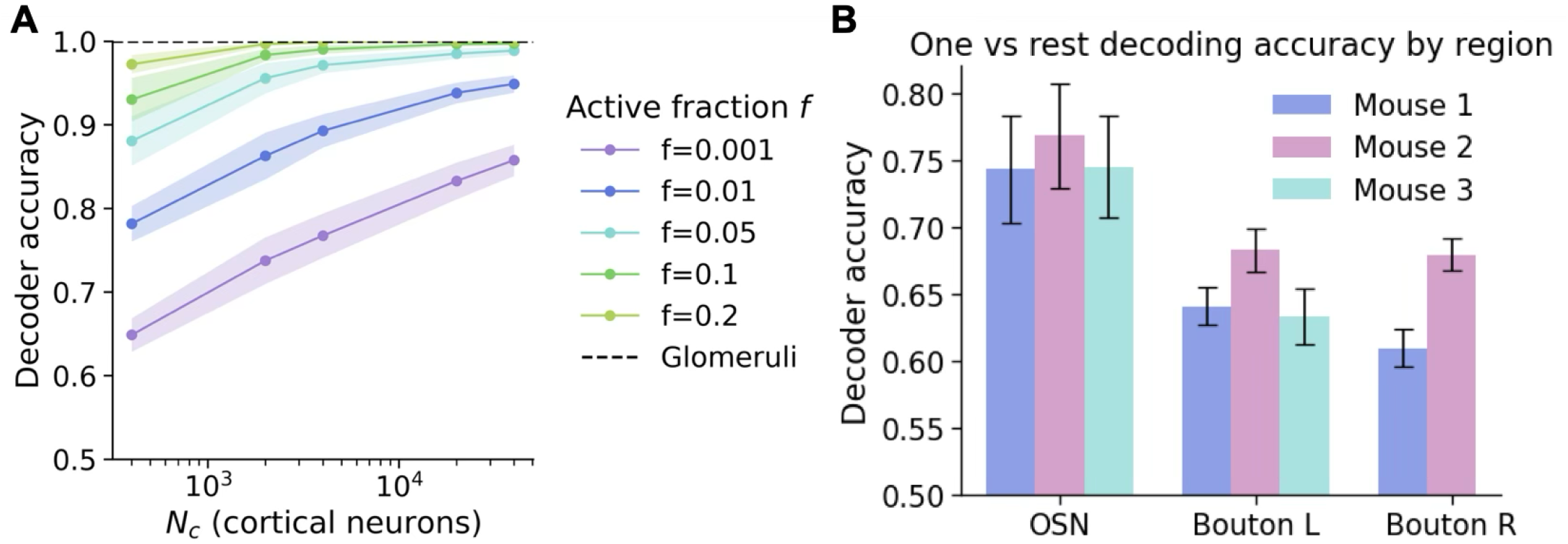
Odor decodability comparison between OSN and piriform cortex. (A) We compare the odor decodability from the glomerular model against a sparse expansive coding model with varying fractions of active neurons and dimensionality. See Methods for simulation details.(B) Zak et al. 2024 recorded population OSN and piriform cortical feedback bouton responses to 100 odor mixtures with complexities 1, 2, 4, 8, 12, and 16 in awake, untrained mice^58^. Boutons were recorded in two different hemispheres (Left and Right, L and R) in two mice. The activity was z-scored before computing the decoding accuracy in the same manner as (A). Error bars represent the mean while error bars denote standard error of the 16 odor classification tasks for each mouse and region.

**Figure S8.**
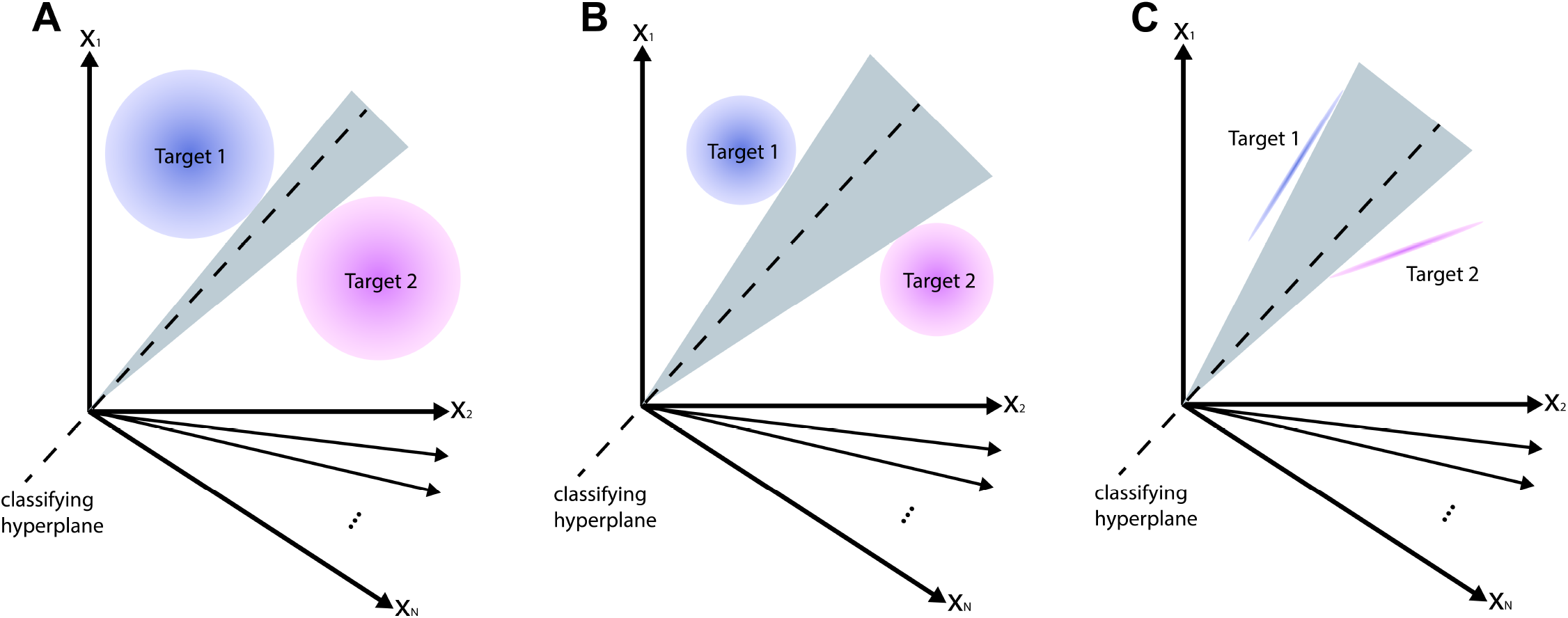
Schematic illustration of manifold capacity and geometry. (**A**) Target odor manifolds are the point cloud of neural response vectors to odor mixtures containing the target odor. Manifolds exist in N-dimensional ambient neural space for N glomeruli but may vary along a lower number of dimensions. In gray, we indicate a cone of valid separating hyperplanes that classify the two target neural manifolds. (**B**) When the magnitude of variability of the manifold decreases, we measure a lower effective radius. This increases the size of the cone of possible separating hyperplanes, which corresponds to a higher manifold capacity. (**C**) When the number of dimensions along which the manifold varies decreases, we measure a lower effective dimensionality. This also increases the size of the cone of possible separating hyperplanes, which corresponds to a higher manifold capacity.

